# Aurora B-dependent Ndc80 Degradation Regulates Kinetochore Composition in Meiosis

**DOI:** 10.1101/836668

**Authors:** Jingxun Chen, Andrew Liao, Emily N Powers, Hanna Liao, Lori A Kohlstaedt, Rena Evans, Ryan M Holly, Jenny Kim Kim, Marko Jovanovic, Elçin Ünal

**Affiliations:** Department of Molecular and Cell Biology, University of California, Berkeley, CA 94720, United States; UC Berkeley QB3 Proteomics Facility, University of California, Berkeley, CA 94720, United States; Fred Hutchinson Cancer Research Center, Seattle, WA 98109, United States; Department of Biology, Columbia University, New York City, NY 10027, United States

**Keywords:** meiosis, kinetochore, Aurora B, Ndc80, chromosome, proteolysis, APC

## Abstract

The kinetochore complex is a conserved machinery that connects chromosomes to spindle microtubules. During meiosis, the kinetochore is restructured to accommodate a specialized chromosome segregation pattern. In budding yeast, meiotic kinetochore remodeling is mediated by the temporal changes in the abundance of a single subunit called Ndc80. We have previously described the regulatory events that control the timely synthesis of Ndc80. Here, we report that Ndc80 turnover is also tightly regulated in meiosis: Ndc80 degradation is active in meiotic prophase, but not in metaphase I. Ndc80 degradation depends on the ubiquitin ligase APC^Ama1^ and is mediated by the proteasome. Importantly, Aurora B-dependent Ndc80 phosphorylation, a mark that has been previously implicated in correcting erroneous microtubule–kinetochore attachments, is essential for Ndc80 degradation in a microtubule-independent manner. The N-terminus of Ndc80, including a 27-residue sequence and Aurora B phosphorylation sites, is both necessary and sufficient for kinetochore protein degradation. Finally, defects in Ndc80 turnover predispose meiotic cells to chromosome mis-segregation. Our study elucidates the mechanism by which meiotic cells modulate their kinetochore composition through regulated Ndc80 degradation, and demonstrates that Aurora B-dependent regulation of kinetochores extends beyond altering microtubule attachments.

## INTRODUCTION

Reproduction is a fundamental feature of life and depends on the accurate segregation of chromosomes from one generation to the next. In eukaryotes, a conserved protein complex known as the kinetochore mediates chromosome segregation. Research over the past three decades has identified at least 40 different proteins that constitute the core of this essential machinery (reviewed extensively in Biggins 2013). While the function of individual kinetochore components has been well established, much less is understood about how the levels of specific subunits are regulated under varying cellular states and how these changes affect kinetochore function.

The kinetochore is composed of two distinct parts: inner and outer kinetochore. The inner kinetochore subunits associate with the chromosome at the centromere, while the outer kinetochore components interact with spindle microtubules. It has been shown that changes in either part can have a profound impact on kinetochore activity and genome inheritance, with potentially deleterious consequences. For example, overexpression of the centromeric histone CENP-A, a component of the inner kinetochore, in yeast, flies, and human cells causes chromosome mis-segregation and genomic instability (Heun et al. 2006; Au et al. 2008; Shrestha et al. 2017). Additionally, overexpression of the outer kinetochore subunits, such as Hec1 (also known as Ndc80) or SKA1, has been observed in many types of cancers and implicated in tumorigenesis (Chen et al. 1997; Hayama et al. 2006; Li et al. 2014; Shen et al. 2016; Chen et al. 2018).

Aside from these pathological states, changes to kinetochore composition also occur in physiological contexts. In various organisms, the kinetochore undergoes extensive remodeling during meiotic differentiation (Asakawa et al. 2005; Sun et al. 2011; Miller et al. 2012; Kim et al. 2013; Meyer et al. 2015), which is the developmental program that generates reproductive cells through two consecutive nuclear divisions. Specifically in budding yeast, the Ndc80 complex disassembles in early meiosis and reassembles during the meiotic divisions, thereby restricting kinetochore activity in a temporal fashion (Asakawa et al. 2005; Miller et al. 2012; Meyer et al. 2015; Chen et al. 2017). This dynamic kinetochore behavior is driven by the fluctuating Ndc80 levels, which are barely detectable in meiotic prophase but become highly abundant during the meiotic divisions. Failure to temporally regulate Ndc80 protein levels and kinetochore activity causes defects in meiotic chromosome segregation and gamete inviability (Miller et al. 2012; Chen et al. 2017), highlighting the importance of Ndc80 regulation.

One way to regulate Ndc80 protein levels occurs through controlling Ndc80 synthesis. Ndc80 production is relatively high during the meiotic divisions, but is completely shut down in meiotic prophase (Figure 1A) (Chen et al. 2017; Chia et al. 2017). This repression in synthesis requires the expression of a meiosis-specific, 5’-extended mRNA expressed from an alternate *NDC80* promoter. This transcript, called LUTI for <u>L</u>ong <u>U</u>ndecoded <u>T</u>ranscript Isoform, is induced by the transcription factor complex Ime1-Ume6 after meiotic entry and cannot be translated into Ndc80 protein. Instead, *NDC80*^*LUTI*^ expression acts to interfere with the transcription of the canonical, protein-coding *NDC80* mRNA isoform. As a result, in meiotic prophase, a stage when *NDC80*^*LUTI*^ is highly expressed, Ndc80 protein synthesis is turned off. After cells exit from meiotic prophase, transcription of the coding *NDC80* isoform is induced by another transcription factor called Ndt80, leading to re-synthesis of Ndc80 and kinetochore activation (Chen et al. 2017). Thus, the developmentally coordinated toggling between these two functionally distinct mRNA isoforms controls Ndc80 production in meiosis.

**Figure 1.**
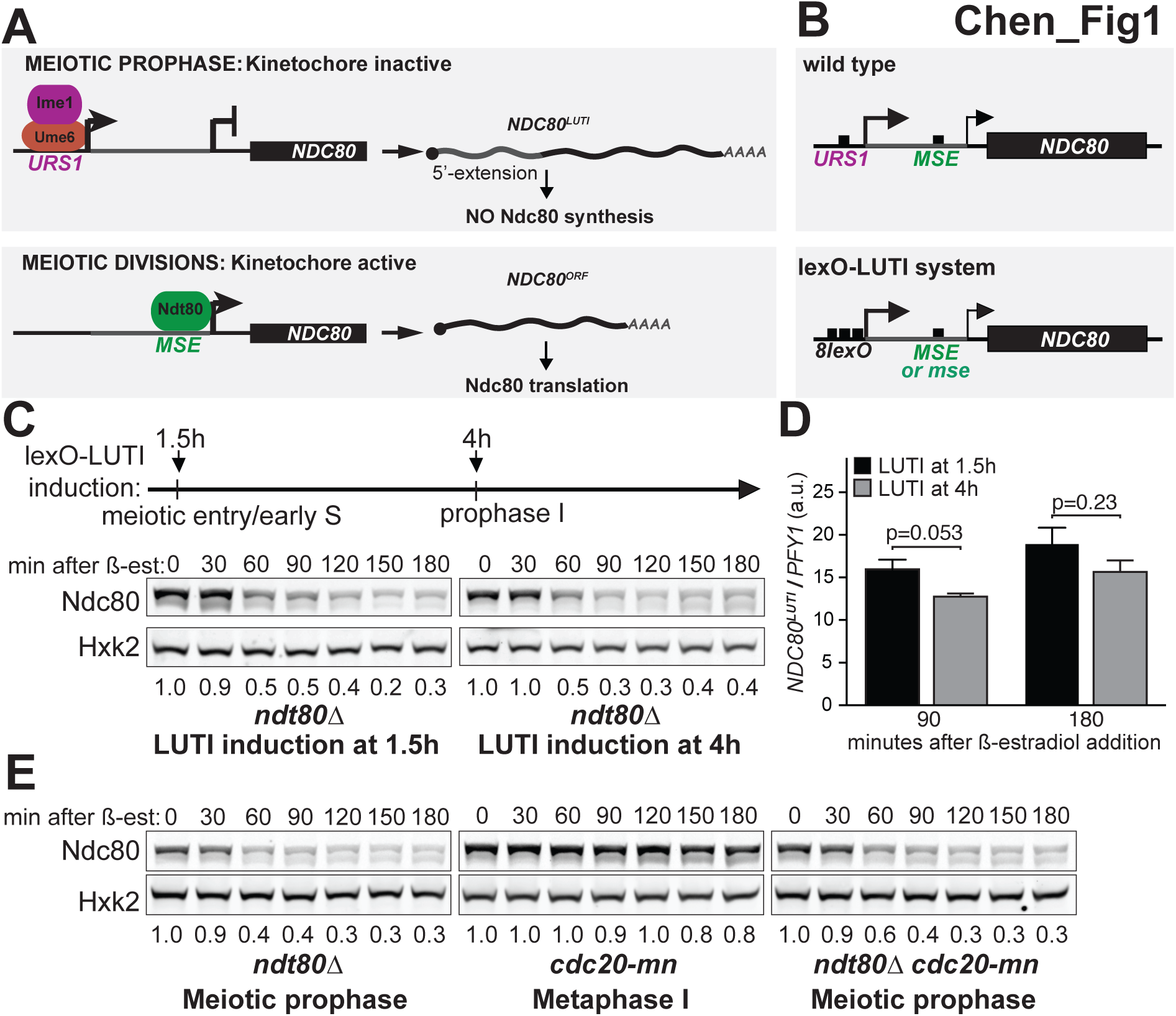
Ndc80 degradation is temporally regulated during meiosis. (A) LUTI-based regulation of Ndc80 protein synthesis in budding yeast meiosis. In meiotic prophase, the Ime1-Ume6 transcription factor complex induces a <u>L</u>ong <u>U</u>ndecoded <u>T</u>ranscript <u>I</u>soform of *NDC80* called *NDC80*^*LUTI*^, which cannot produce Ndc80 protein due to the upstream open reading frames in its 5’ extension. *NDC80*^*LUTI*^ represses transcription of a protein-coding isoform of *NDC80*, *NDC80*^*ORF*^. Through this combined act of transcriptional and translational repression, *NDC80*^*LUTI*^ inhibits Ndc80 protein synthesis. In the meiotic divisions, *NDC80*^*ORF*^ is induced by a second meiotic transcription factor, Ndt80. URS1 (upstream regulatory sequence 1), a DNA binding motif for Ume6. MSE (mid-sporulation element), a DNA binding motif for Ndt80. (B) The lexO-LUTI system induces *NDC80*^*LUTI*^ expression upon ß-estradiol addition, thus conditionally inhibiting *NDC80*^*ORF*^ expression and Ndc80 protein synthesis. Top: Regulatory elements of the *NDC80* gene. Bottom: The lexO-LUTI system. *mse*, a mutant *MSE* site defective in Ndt80 binding. (C) Ndc80 turnover in early or late meiotic prophase. The strain carrying the lexO-LUTI (UB14883) was transferred to the sporulation medium (SPO) at 0 h to induce meiosis, and ß-estradiol was added at either 1.5 h or at 4 h after meiosis induction. The strain was halted in meiotic prophase using an *ndt80∆* block. Here and throughout, Ndc80 levels were determined by anti-V5 immunoblot. Hxk2, loading control. Unless specified, the numbers below the immunoblots were calculated by first normalizing Ndc80 levels to Hxk2 levels in each lane, and then dividing the ratio to the 0 h time point. All the experiments in this study were performed at least two times, and one representative biological replicate is shown. (D) Induction levels of *NDC80*^*LUTI*^ mRNA for the experiment in panel C, measured by reverse transcription followed by quantitative PCR (RT-qPCR). For all RT-qPCR experiments, *NDC80*^*LUTI*^ signals were normalized to that of *PFY1*. a.u., arbitrary unit. The mean from 3 independent experiments, along with the standard error of the mean, is displayed. The p-values were calculated by a two-tailed student’s t-test. (E) Ndc80 turnover in late meiotic prophase or in metaphase I arrest. The *ndt80∆* (UB19616) and the *ndt80∆ cdc20-mn* (UB19618) strains were cultured in SPO for 4 h before ß-estradiol addition. Both strains were halted in meiotic prophase with an *ndt80∆* block. The *cdc20*-meiotic null mutant (*cdc20-mn*, UB19678) was cultured in SPO for 5 h before ß-estradiol addition and subsequently halted in metaphase I for 3 h.

The LUTI-based regulation explains how meiotic cells can effectively repress Ndc80 protein synthesis. However, since Ndc80 is clearly detected at meiotic entry (Asakawa et al. 2005; Miller et al. 2012; Meyer et al. 2015; Chen et al. 2017), regulated Ndc80 synthesis alone cannot fully explain kinetochore inactivation in meiotic prophase. Additional mechanisms must be in place to clear the existing pool of Ndc80 such that the kinetochores can disassemble in a timely manner. Interestingly, the human homolog of Ndc80, Hec1, undergoes degradation in a cell cycle-dependent manner, but the turnover mechanism remains elusive (Ferretti et al. 2010). More generally, little is known about the factors that mediate kinetochore subunit degradation in a developmental context.

Here, we describe the mechanism by which Ndc80 degradation is controlled in budding yeast meiosis. We found that the degradation of Ndc80 is temporally regulated. Its proteolysis in meiotic prophase requires Aurora B/Ipl1 kinase-dependent phosphorylation, which has been previously linked to correcting erroneous microtubule– kinetochore attachments (reviewed in Biggins 2013). The N-terminus of Ndc80, including a 27-residue sequence and Ipl1 phosphorylation sites, is both necessary and sufficient for kinetochore protein degradation. In addition to phosphorylation, Ndc80 degradation depends on the ubiquitin ligase APC^Ama1^ and proteasome activity. Failure to degrade Ndc80 causes premature kinetochore assembly in meiotic prophase and predisposes cells to meiotic chromosome segregation defects. Our results provide mechanistic insight into how cells can developmentally modulate kinetochore composition through subunit proteolysis and highlight the importance of timely Ndc80 degradation in promoting accurate meiotic chromosome segregation.

## RESULTS

### Ndc80 degradation is temporally regulated in meiosis

In meiotic prophase, the residual Ndc80 protein from the pre-meiotic cell cycle is turned over by an unknown mechanism (Chen et al. 2017). To study Ndc80 degradation without a confounding effect from its synthesis regulation, we took advantage of a previously established method, which allowed us to turn off Ndc80 synthesis in a conditional manner (Chia et al. 2017). Specifically, we used a strain in which the endogenous *NDC80*^*LUTI*^ promoter was replaced with an inducible promoter controlled by an array of 8 lex operators (*8lexO*) (Figure 1B). The same strain carries a chimeric lexA-B112 transcription factor fused to an estradiol-binding domain (lexA-B112-ER), which allows inducible transcription from the *8lexO* promoter in the presence of ß-estradiol (Ottoz et al. 2014). Without ß-estradiol (uninduced), the coding *NDC80* transcript (hereafter referred to as *NDC80*^*ORF*^) is expressed and Ndc80 is synthesized. After ß-estradiol addition, *NDC80*^*LUTI*^ is expressed, resulting in repression of Ndc80 synthesis In comparison to wild type cells, this induction system led to similar kinetics of Ndc80 degradation following meiotic entry (Figure S1A).

Using this system, we examined Ndc80 turnover at different stages of meiosis to determine the specific time window of Ndc80 degradation. We treated cells with ß-estradiol either close to meiotic entry (1.5h after meiotic induction) or later (4h after meiotic induction). Meanwhile, the cells were held in meiotic prophase by deletion of *NDT80*, which encodes a transcription factor required for meiotic progression. We found that Ndc80 was degraded with similar kinetics in either condition (Figure 1C). The levels of *NDC80*^*LUTI*^ induction were also similar, as measured by reverse transcription followed by quantitative polymerase chain reaction (RT-qPCR) (Figure 1D), suggesting that Ndc80 synthesis was successfully repressed. This result suggests that Ndc80 turnover can occur throughout meiotic prophase.

To determine if Ndc80 is degraded beyond meiotic prophase, we monitored Ndc80 levels during a metaphase I arrest induced by Cdc20 depletion (*cdc20-mn*). Cdc20 is an activator of the anaphase-promoting complex, APC/C, a ubiquitin ligase necessary for metaphase-to-anaphase transition (Visintin et al. 1997; Hwang et al. 1998; Yu 2007). For this experiment, we mutated the Ndt80 binding site (also known as <u>m</u>id <u>s</u>porulation <u>e</u>lement or MSE) at the *NDC80* promoter (Chen et al. 2017). This alteration is required because the second burst of Ndc80 synthesis, which depends on the MSE site, occurs after cells exit meiotic prophase. Mutating this site ensures that Ndc80 synthesis can be repressed by ß-estradiol addition even after meiotic prophase. We found that while Ndc80 was degraded in meiotic prophase, it remained remarkably stable during the metaphase I arrest induced by *cdc20-mn* (Figure 1E). The level of *NDC80*^*LUTI*^ induction was ~40% lower in *cdc20-mn* cells than in wild type (Figure S1B). In principle, this reduction of *NDC80*^*LUTI*^ could cause an increase in Ndc80 synthesis, leading to higher protein levels. To exclude this possibility, we used cycloheximide to globally inhibit protein synthesis. Ndc80 was still stable during the metaphase I arrest and degraded in late prophase I under these conditions (Figure S1C), suggesting that the stability of Ndc80 protein differed between the two states. While it is possible that APC^Cdc20^ may regulate Ndc80 degradation in metaphase I, we found that Cdc20 was dispensable for Ndc80 degradation in meiotic prophase (*ndt80∆*, *cdc20-mn*) (Figure 1E). We conclude that Ndc80 degradation is temporally regulated, occurring in a meiotic prophase-specific manner.

### Proteasome activity and Aurora B/Ipl1 regulate Ndc80 degradation

To identify the regulators of Ndc80 degradation, we surveyed the key cellular and proteolytic events in meiotic prophase. We found that synapsis, recombination, and DNA replication were all dispensable for Ndc80 degradation. Ndc80 degradation was normal in *spo11∆* cells despite the lack of recombination and synapsis (Giroux et al. 1989; Cao et al. 1990; Keeney et al. 1997) (Figure 2A). Ndc80 degradation also occurred normally in cells depleted of the DNA replication factor Cdc6 (*cdc6-mn*) (Hochwagen et al. 2005; Brar et al. 2009; Blitzblau et al. 2012), consistent with a previous report (Meyer et al. 2015) (Figure 2A). These results suggest that Ndc80 degradation, and thus kinetochore remodeling, is independent of major meiosis-specific changes to chromosomes.

**Figure 2.**
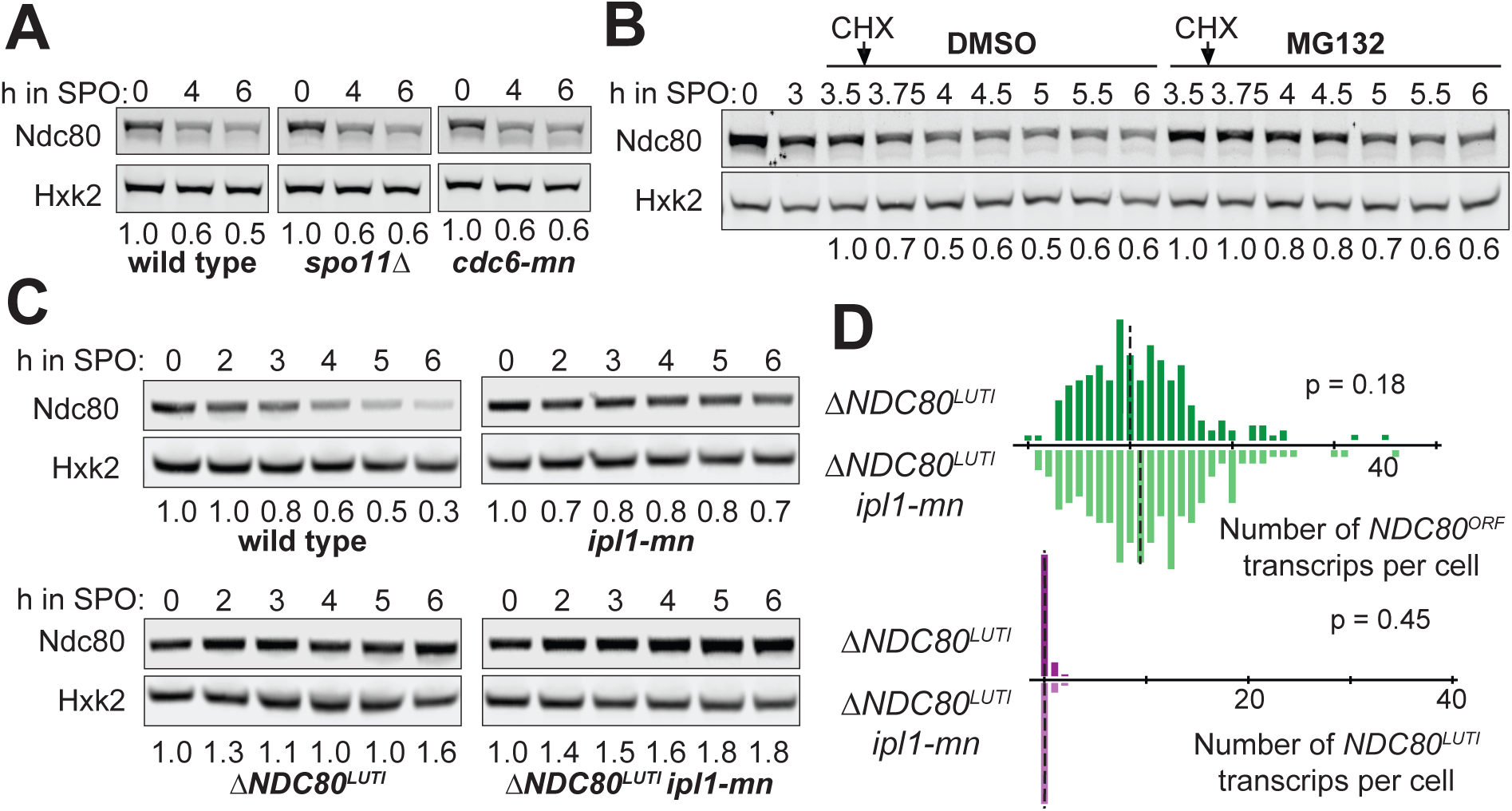
Proteasome and the Aurora B/Ipl1 kinase regulate Ndc80 degradation. (A) Dependency of Ndc80 protein degradation on recombination, synapsis, or DNA replication. The wild type (UB1338), *spo11∆* (UB11793), and *cdc6-*meiotic null (*cdc6-mn*, UB13656) strains were transferred to SPO at 0 h and halted in meiotic prophase using the *pGAL-NDT80 GAL4.ER* system until 6 h in SPO. (B) Dependency of Ndc80 degradation on active proteasomes. Cells (*pdr5∆*, UB2405) were induced to sporulate at 0 h. After 3 h in SPO, cells were split and treated with either DMSO or the proteasome inhibitor MG132 (100 µM). 30 min later, cycloheximide (CHX) was added (0.2 mg/ml). For either condition (DMSO or MG132), all the time points were normalized to the time point immediately before the cycloheximide addition (3.5 h). (C) The effects on Ndc80 levels when the meiotic depletion of *IPL1* (*ipl1-mn*) was combined with a mutant that fails to repress Ndc80 synthesis (*∆NDC80*^*LUTI*^). The wild type (UB1338), *ipl1-mn* (UB1013), *∆NDC80*^*LUTI*^ (UB2932), and *∆NDC80*^*LUTI*^ *ipl1-mn* (UB3948) cells were sporulated as in (A). (D) smFISH quantification of the *NDC80*^*ORF*^ and *NDC80*^*LUTI*^ mRNA levels in the *∆NDC80*^*LUTI*^ cells (UB2932) and *∆NDC80*^*LUTI*^ *ipl1-mn* (UB3948) cells in meiotic prophase. Samples were taken at 5 h in SPO. The relative frequency histograms of the cells with a given number of *NDC80*^*LUTI*^ and *NDC80*^*ORF*^ transcripts per cell were graphed using the data pooled from 2 independent experiments. Dashed line, the median number of transcripts per cell. A total of 218 cells were counted for *∆NDC80*^*LUTI*^, and 208 cells, for *∆NDC80^LUTI^ ipl1-mn*. Two-tailed Wilcoxon Rank Sum test was performed.

Next, we tested the ubiquitin-proteasome system, a key protein degradation pathway in the cell (Finley et al. 2012). In meiotic prophase, proteasomes are localized to chromosomes (Ahuja et al. 2017) and could mediate Ndc80 degradation. We treated meiotic prophase cells with the proteasome inhibitor MG132. Compared to the vehicle control, Ndc80 levels were modestly stabilized within 1h of MG132 treatment (Figure S2A). During this time, the expression of *NDC80*^*LUTI*^ did not change significantly, but it had decreased by ~50% by the second hour of treatment (Figure S2B). To exclude the confounding effects from the differences in Ndc80 synthesis, we performed a cycloheximide-chase experiment. The levels of Ndc80 were stabilized in the first hour of cycloheximide treatment although they eventually declined (Figure 2B). This result indicates that the proteasome contributes, at least partially, to Ndc80 degradation.

Previously, it has been shown that the decline of Ndc80 levels in meiotic prophase requires the kinase Aurora B/Ipl1 (Meyer et al., 2015), raising the possibility that Ipl1 regulates Ndc80 degradation. We confirmed that Ndc80 abundance was increased upon meiotic depletion of Ipl1 (*ipl1-mn*) (Figure 2C). This increase was not the result of elevated Ndc80 synthesis, as shown by three observations. First, the level of *NDC80*^*LUTI*^ mRNA was not altered in *ipl1-mn* mutants (Figure S2C). Second, Ipl1 depletion increased Ndc80 levels additively with a mutant that fails to repress Ndc80 synthesis (*∆NDC80*^*LUTI*^) (Figure 2C). Finally, the expression of the protein-coding *NDC80*^*ORF*^ isoform did not significantly change in the double mutant (*∆NDC80*^*LUTI*^ *ipl1-mn*) compared to the single mutant (*∆NDC80*^*LUTI*^), as shown by single molecule RNA fluorescence *in situ* hybridization (smFISH) (Figure 2D and S2D). Based on these data, we conclude that Ipl1 regulates Ndc80 turnover rather than synthesis.

### Ndc80 degradation requires Ipl1-mediated phosphorylation

How does Ipl1 regulate Ndc80 abundance mechanistically? In one model, Ipl1 depletion may alter Ndc80 abundance indirectly by affecting microtubule behavior in meiotic prophase. At this meiotic stage, the yeast centrosomes, known as the spindle pole bodies (SPBs), are duplicated but prevented from separating to form spindle microtubules. Meanwhile, the kinetochores become dispersed from the SPBs, presumably due to the lack of microtubule-kinetochore interactions (Kim et al. 2013; Meyer et al. 2013; Meyer et al. 2015). Both kinetochore dispersion and inhibition of spindle formation in meiotic prophase require Aurora B/Ipl1 activity (Kim et al. 2013; Meyer et al. 2013; Meyer et al. 2015). *ipl1-mn* mutants prematurely separate the duplicated SPBs and form long microtubules (Shirk et al. 2011; Kim et al. 2013). These microtubules interact with the kinetochores, leading to kinetochore re-clustering (Meyer et al. 2013; Meyer et al. 2015). In this context, microtubules could shield Ndc80 from degradation. Accordingly, the effect of *ipl1-mn* would depend on the presence of microtubules. However, when we depolymerized microtubules in *ipl1-mn* cells using a microtubule poison cocktail (benomyl & nocodazole), Ndc80 remained stable in meiotic prophase (Figure S3A-C). Thus, it is unlikely that Ipl1 indirectly regulates Ndc80 degradation by affecting microtubule behavior.

Since Ipl1 is a kinase, it may promote Ndc80 turnover by phosphorylating factors that turn over Ndc80, or by phosphorylation Ndc80 itself. Previous work has shown that Ndc80 is a direct substrate of Ipl1 (Cheeseman et al. 2002; Akiyoshi et al. 2009). Ndc80 has seven known Ipl1 consensus sites ([KR]-X-[ST]-[ILVST]) (Cheeseman et al. 2002), which are required for Ipl1 to phosphorylate Ndc80 *in vitro* (Akiyoshi et al. 2009). To test whether Ndc80 is degraded through a phosphorylation-dependent mechanism, we first asked if Ndc80 is phosphorylated at the time of its degradation. We immunoprecipitated Ndc80 from meiotic prophase cells 1h after MG132 treatment and analyzed the post-translational modifications by mass spectrometry. We found two recurring, high confidence phosphorylation sites T54 and T248 on Ndc80 (Figure 3A). The first site has been previously characterized and contains an Ipl1 consensus site (Cheeseman et al. 2002; Akiyoshi et al. 2009), whereas the second site lacks the Ipl1 consensus sequence.

**Figure 3.**
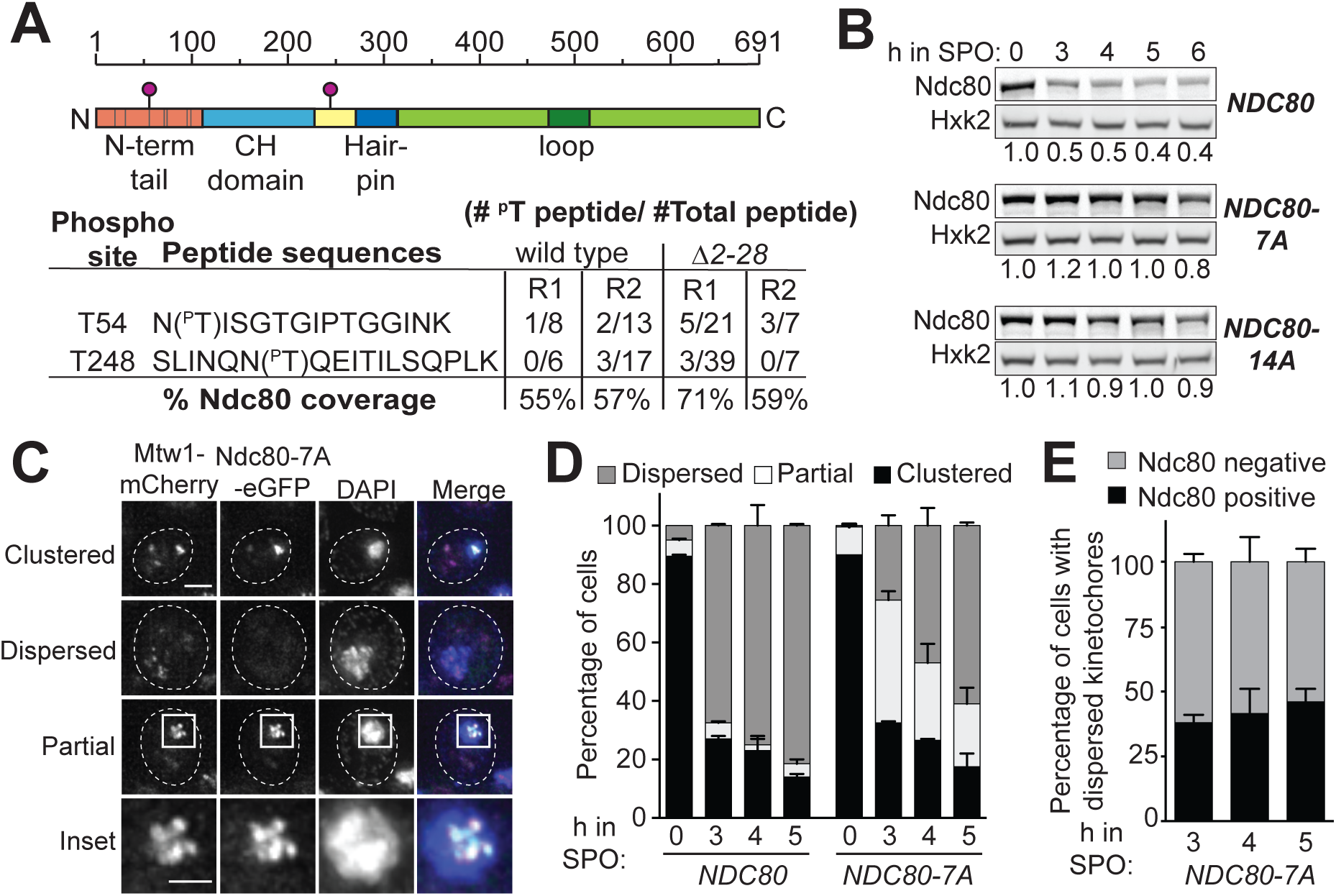
Ndc80 degradation requires Ipl1-mediated phosphorylation. (A) Phosphorylation sites detected on wild type Ndc80 proteins and Ndc80(∆2-28) proteins in meiotic prophase by mass spectrometry (MS). The wild type (*pdr5∆*, UB2405) cells were treated with MG132 at 3 h after transfer to SPO, and samples were collected 1 h after treatment. For the *∆2-28* cells (UB5662), samples were collected after 5 h in SPO. Top: schematic of Ndc80 and the *in vivo* phosphorylation sites detected by MS (pink circles). Gray bars, the 7 Ipl1 consensus sites. N-term, N terminal. CH, calponin homology. Bottom: Detected Ndc80 phosphopeptides. The detected number of phosphopeptides and total peptides (phosphorylated and unmodified combined), as well as the overall sequence coverage of Ndc80, are reported for two biological replicates. R1, repeat 1; R2, repeat 2. (B) Ndc80 protein level in meiotic prophase for the wild type (UB4074), *NDC80-7A* (UB13658), *NDC80-14A* (UB17707) cells. Samples were taken and processed as in Figure 2A. (C) Representative images of the clustered, dispersed, and partially clustered kinetochores in *NDC80-7A* (UB15701) cells, which were fixed at 5 h after transfer to SPO. Mtw1 was tagged with mCherry and Ndc80-7A tagged with eGFP. DNA was stained with DAPI. Scale bar: 2 µm. Inset scale bar: 1 µm. (D) The percentage of cells with clustered, partially clustered, or dispersed kinetochores in meiotic prophase. The wild type (UB1083) and *NDC80-7A* (UB15701) cells were fixed immediately after transfer to SPO (0 h) and at 3 h, 4 h, and 5 h later. 100 cells were counted per time point. The mean and the range of the percentage for two biological replicates are graphed. (E)The percentage of cells with dispersed kinetochores that contained Ndc80-7A-eGFP signal on at least one kinetochore at the indicated time points.

Next, we mutated the serine or threonine in all seven Ipl1 consensus sites, generating the allele (*NDC80-7A*) known to greatly reduce Ndc80 phosphorylation by Ipl1 *in vitro* (Akiyoshi et al. 2009). The mutations did not affect *NDC80*^*LUTI*^ expression (Figure S3D), ruling out the possibility that the synthesis repression of Ndc80 was disrupted. We found that Ndc80-7A was highly stable in meiotic prophase (Figure 3B). Mutating additional serine and threonine residues around T54 and T248 (*NDC80-14A*) did not enhance Ndc80 stabilization (Figure 3B). Thus, we conclude that the seven Ipl1 consensus sites at the N-terminal region of Ndc80 are required for its turnover in meiotic prophase.

We further asked if the stabilized Ndc80-7A proteins localized to the kinetochores and affected kinetochore behavior. We fused Ndc80-7A to the enhanced green fluorescent protein (eGFP), and generated a mCherry-tagged allele for the inner kinetochore protein Mtw1, which remains chromosome-associated throughout meiotic prophase. For both the wild type and *NDC80-7A* mutant, over 85% of the cells had clustered kinetochores at meiotic entry (0h in SPO), and all of the kinetochore clusters had Ndc80-eGFP signal (Figure 3C-D). When the wild type cells progressed into meiotic prophase (3-5h in SPO), the kinetochores became dispersed in >70% of the cells, and none of the dispersed kinetochores had Ndc80-eGFP signal (Figure 3D). In contrast, fewer than 30% of the *NDC80-7A* cells had dispersed kinetochores after 3h in SPO (Figure 3D). Instead, ~40% of the cells had partially clustered kinetochores (Figure 3C, partial and inset), all of which had Ndc80-7A-eGFP on them. By 5h, >50% of the *NDC80-7A* cells had dispersed kinetochores (Figure 3D), and about half of these cells had Ndc80-7A-eGFP on at least one kinetochore (Figure 3E). These results demonstrate that the stabilized Ndc80-7A proteins localize to the kinetochore and affect the timing of kinetochore dispersion in meiotic prophase.

### Ndc80 degradation requires a specific sequence at its N-terminus

Since the seven Ipl1 consensus sites are located in the 112-residue N-terminal tail of Ndc80, we systematically truncated this region to narrow down the sites necessary for Ndc80 degradation. We found residues 2-28 to be necessary for the decline of Ndc80 levels (Figure 4A, top panel). Within this segment, residues 11-19 were the most critical (Figure 4A, bottom panel). To our surprise, the decline of Ndc80 levels was not altered when the 11 serines and threonines in the first 30 residues of Ndc80 were mutated (underlined in Figure 4B) (*NDC80-11A*), indicating that Ndc80 degradation does not depend on the potential phosphorylation sites within this region. Ndc80 turnover was also normal when the 4 histidines in the 11-19 region (green letters in Figure 4B) were mutated to either alanines (*NDC80-4A*) or leucines (*NDC80-4L*) (Figure S4A).

**Figure 4.**
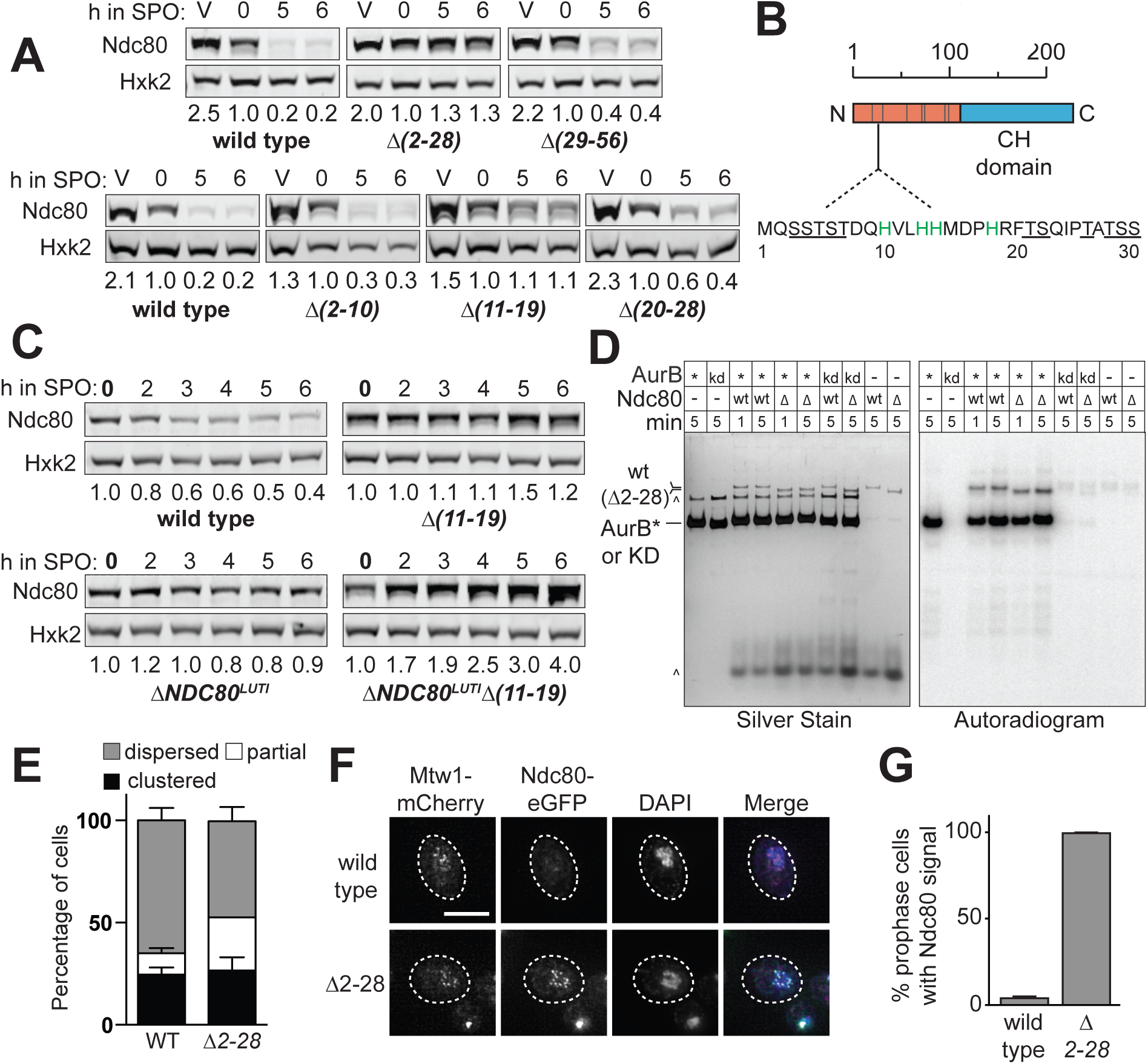
Ndc80 degradation requires a short sequence at its N terminus. (A) Truncation analysis of Ndc80 to identify the residues necessary for Ndc80 degradation. Strains harboring the deletions of residues 2-28 *(∆2-28*, UB5662), residues 29-56 (*∆29-56*, UB15972), residues 2-10 (*∆2-10*, UB7039), residues 11-19 (*∆11-19*, UB7029), and residues 20-28 (*∆20-28*, UB7031) were sporulated along with wild type cells (UB4074) as in Figure 2A. The vegetative samples (V) were taken while each strain was growing exponentially in rich medium. Note: Hxk2 level declined slightly as meiotic prophase progressed. Thus, the normalized level of Ndc80 became over 1.0 in the later stages of meiotic prophase. (B) An abridged schematic of Ndc80. The sequence of the first 30 residues is displayed. Gray bars, the 7 Ipl1 consensus sites. The underlined residues are the 11 serines and threonines mutated in the *11A* mutant, and the green residues are the 4 histidines mutated in the *4(H to A)* and *4(H to L)* mutants. (C) The effects on Ndc80 levels when the *∆11-19* mutant was combined with a mutant that fails to repress Ndc80 synthesis (*∆NDC80*^*LUTI*^). The wild type (UB4074), *∆11-19* (UB7029), *∆NDC80*^*LUTI*^ (UB11797), and *∆NDC80*^*LUTI*^ *∆11-19* (UB11799) cells were sporulated as described in Figure 2A. (D) *In vitro* kinase assay for the wild type Ndc80 and Ndc80(∆2-28) proteins purified from UB16284 and UB19957, respectively. The Ndc80 proteins were phosphorylated *in vitro* by either 1 µM recombinant AurB* (fusion of the C-terminal activation box of INCEP^sli15^ to Aurora B^Ipl1^) or 1 µM kinase-dead AurB* (KD) and γ-^32^P-ATP. All reactions were performed at room temperature for either 1 min or 5 min, analyzed by SDS-PAGE, and then visualized by silver staining and autoradiography. wt, wild type Ndc80. ∆, Ndc80(∆2-28). *, AurB*. ^, background bands. (E) The percentage of the wild type or *∆2-28* cells with dispersed, partially clustered, and clustered kinetochores after 3 h in SPO. The average, as well as the range, of 2 independent biological replicates are displayed. Over 100 meiotic cells were counted. (F) Representative images of the wild type Ndc80 and Ndc80(∆2-28) proteins in meiotic prophase. The wild type (UB1083) and *∆2-28* (UB15619) cells were fixed after 5 h in SPO. Mtw1 was tagged with mCherry and Ndc80(∆2-28) tagged with eGFP. DNA was stained with DAPI. Scale bar: 5 µm. (G) The average percentages of meiotic prophase cells (identified by the pachytene DAPI morphology) with colocalized Ndc80-eGFP and Mtw1-mCherry signals, as well as the range, of 2 independent biological replicates are displayed. At least 70 meiotic prophase cells were counted.

How does the 2-28 region regulate Ndc80 abundance? We first confirmed that the 2-28 region regulates Ndc80 degradation rather than Ndc80 synthesis. The wild type, *∆2-28* and *∆11-19* cells had similar *NDC80*^*LUTI*^ levels in meiotic prophase (Figure S4B). Also, the Ndc80 protein levels were additively increased when the *∆11-19* mutant was combined with the mutant that fails to inhibit Ndc80 synthesis (*∆NDC80*^*LUTI*^) (Figure 4C). Despite the difference in protein levels, the expression of the coding *NDC80*^*ORF*^ mRNA was not significantly different between the double mutant (*∆NDC80^LUTI^ ∆11-19*) and the single mutant (*∆NDC80*^*LUTI*^)(Figure S4C-D), suggesting that the 2-28 residues regulate Ndc80 stability.

Since Ndc80 degradation requires Ipl1-dependent phosphorylation, we next asked if such phosphorylation depends on the 2-28 residues. We immunoprecipitated Ndc80(∆2-28) protein from meiotic prophase and performed mass spectrometry. Both phosphorylation sites (T54 and T248) observed in the wild type Ndc80 protein were detected (Figure 3A). In addition, we performed an *in vitro* kinase assay using a recombinant Ipl1 protein variant (AurB*) (de Regt et al. 2018) and found no noticeable difference in the degree of Ndc80 phosphorylation between the wild type and Ndc80(∆2-28) protein (Figure 4D). These results strongly suggest that the Ipl1-depedent Ndc80 phosphorylation occurs normally in the *∆2-28* mutant. Thus, we conclude that the 2-28 region is required for Ndc80 degradation at a step downstream or in parallel to Ipl1-dependent phosphorylation.

We attempted to understand the role of the 2-28 residues in Ndc80 degradation by looking for changes in the binding partners of the wild type or Ndc80(∆2-28) protein during meiotic prophase. By quantitative mass spectrometry using tandem mass tags, we found that Sis1, a J-domain protein that regulates heat shock protein activity (Kampinga and Craig 2010), interacted with the wild type but not the Ndc80(∆2-28) protein (Figure S4E). However, Sis1 did not appear to be required for Ndc80 degradation since even the mutants that displayed normal Ndc80 degradation (e.g. *∆29-56*) were still defective in Sis1 binding (Figure S4F). It remains unclear how the 2-28 segment mediates Ndc80 degradation.

Despite acting through an unknown mechanism, the 2-28 residues of Ndc80 are evidently important for clearing Ndc80 from the kinetochores and facilitating kinetochore dispersion in meiotic prophase. For the *∆2-28* mutant, we observed an increased proportion of cells with partially clustered kinetochores after 3h in SPO (~25%, Figure 4E), although the percentage was lower than that of the *NDC80-7A* mutant (~40%, Figure 3D). By 5h in SPO, most of the *∆2-28* cells had dispersed kinetochores in meiotic prophase, and Ndc80(∆2-28)-eGFP signal was detected on all the dispersed kinetochores (Figure 4F-G), demonstrating that the stabilized Ndc80(∆2-28) protein localize to the kinetochore. In addition, our observations suggest that while Ndc80 degradation is not required for kinetochore dispersion, it can affect dispersion kinetics (given the accumulation of cells with partially clustered kinetochores), potentially by acting in conjunction with Ipl1-dependent phosphorylation.

### The N-terminus of Ndc80 is sufficient to induce proteolysis

Both the 2-28 region of Ndc80 and Ipl1-mediated phosphorylation are required for Ndc80 degradation. Are these two features sufficient to induce protein degradation in meiotic prophase? We tested this idea by fusing the N-terminal region of Ndc80 (residues 2-112), including both the 2-28 region and the Ipl1 consensus sites, to an inner kinetochore protein, Ame1 (Figure 5A). The protein levels of Ame1 were stable in meiotic prophase, as shown by a cycloheximide-chase experiment (Figure S5A). We placed this Ame1-tail construct under the promoter of *NDC80*, which leads to synthesis repression in meiotic prophase. Addition of the N-terminal tail of Ndc80 caused Ame1 to become unstable in meiotic prophase in an Ipl1-dependent manner (Figure 5B). Therefore, the N-terminal tail of Ndc80 is sufficient to induce proteolysis in meiotic prophase.

**Figure 5.**
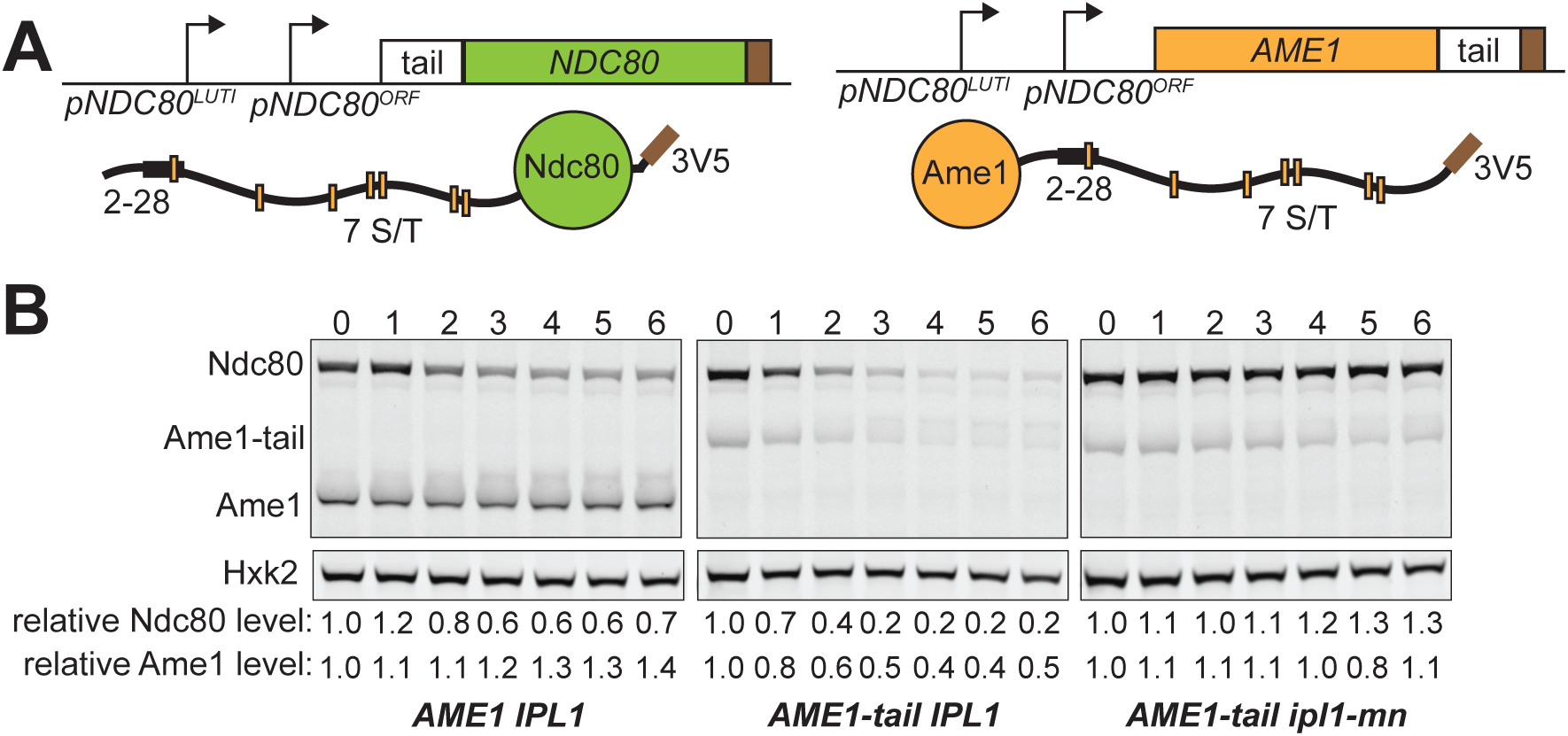
The N-terminus of Ndc80 is sufficient to induce proteolysis. (A) Schematic of the Ndc80 tail-Ame1 fusion. The tail of Ndc80 (residue 2-112), including both the degron sequence and the Ipl1 phosphorylation sites, is fused to the C-terminus of Ame1. The fusion construct is controlled by the *NDC80* promoter, which allows synthesis repression in meiotic prophase. (B) Ame1-tail is degraded in meiotic prophase in an Ipl1-dependent manner. The *IPL1 AME1* (UB20358), *IPL1 AME1-tail* (UB20354), and *ipl1-mn AME1-tail* (UB22397) cells were sporulated as described in Figure 2A. In all the strains, Ndc80 was tagged with 3V5, the same epitope tag for Ame1, as an internal control. The relative Ndc80 levels were calculated as described in Figure 1C. The relative Ame1 levels were calculated by normalizing Ame1 levels to Hxk2 levels in each lane, and then divide the normalized values to that of time 0 h.

### APC^Ama1^ regulates Ndc80 degradation

Since proteasome activity contributes to Ndc80 degradation, we posited that Ndc80 is degraded via a system mediated by ubiquitin/proteasome, which requires one or more E3 ligases to ubiquitinate the substrate for proteasome recognition (Finley et al. 2012). Thus, we surveyed a candidate list of E3 ubiquitin ligases for their roles in Ndc80 degradation. We found the meiosis-specific activator of the APC/C, Ama1, to be required for Ndc80 turnover (Figure 6A). APC^Ama1^ regulates Ndc80 degradation rather than synthesis, because the levels of *NDC80*^*LUTI*^ were not significantly altered in the *ama1∆* mutant (Figure S6A). A cycloheximide-chase experiment further supported that APC^Ama1^ acts at a post-translational step in regulating Ndc80 levels (Figure 6C-D). In addition to Ndc80, the Ame1-tail fusion was also stabilized in an APC^Ama1^-dependent manner (Figure 6B).

**Figure 6.**
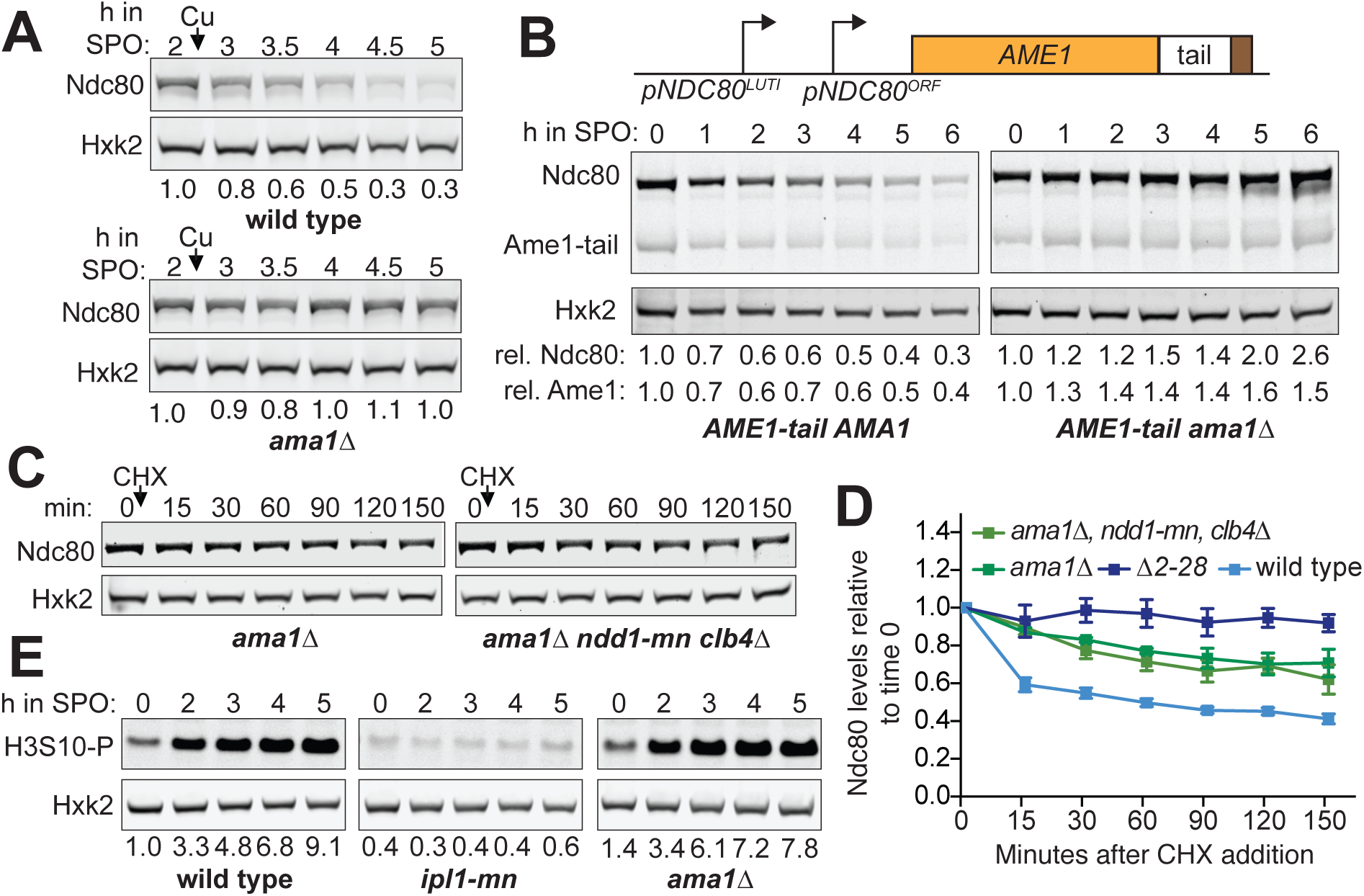
APC^Ama1^ regulates Ndc80 degradation. (A) Dependency of Ndc80 levels on Ama1 in meiotic prophase. Sporulation of the wild type (UB6598) and *ama1∆* (UB22499) cells was synchronized with the *pCUP-IME1/pCUP-IME4* system. Meiotic entry was induced by CuSO_4_ addition after cells were incubated in SPO for 2 h. (B) Dependency of Ame1-tail fusion levels on Ama1 in meiotic prophase. The *AME1-tail AMA1* (UB20354) and *AME1-tail ama1∆* (UB23001) strains were sporulated as in Figure 2A. Quantification of Ndc80 and Ame1 were performed as in Figure 5B. rel, relative. (C) Assessing the requirement of Ndd1 and Clb4 in the Ndc80 stabilization that was induced by *ama1∆*. The *ama1∆* (UB20223), and *pCLB2-NDD1 (ndd1-mn) clb4∆ ama1∆* (UB22850) cells were cultured in SPO for 3 h before a final concentration of 0.2 mg/ml cycloheximide was added. During the entire CHX experiment, both strains were halted in meiotic prophase. (D) The Ndc80 protein levels relative to the time point immediately before CHX addition. The average and the standard errors of the mean for 5 independent experiments are graphed. (G) Ipl1 activity in the *ama1∆* mutant. The wild type (UB3954), *ipl1-mn* (UB1013), and *ama1∆* (UB20223) cells were sporulated as in Figure 2A. The levels of H3 serine 10 phosphorylation (H3S10-P) were detected by a phospho-specific antibody. The number below each lane was first normalized to Hxk2 levels and then to the 0 h time point of the wild type strain.

It has been previously shown that Ama1 deletion causes a mitotic-like cellular state in meiotic prophase (Okaz et al. 2012). This occurs due to stabilization of two APC^Ama1^ substrates: Ndd1, a transcription factor required for the mitotic expression of B-type cyclins and Clb4, a B-type cyclin itself. Therefore, it is possible that such a mitotic-like state in the *ama1∆* mutant could mimic a condition in which Ndc80 is stable, such as metaphase I. However, we found no evidence in support of this model. In our strains, premature spindle formation, a phenotype associated with elevated cyclin/CDK activity, was not apparent in the *ama1∆* mutant (Figure S6B). As an additional test, we removed Ndd1 and Clb4 from the *ama1∆* mutant and found that Ndc80 was still stable (Figure 6C-D, S6C), suggesting that Ama1 deletion does not stabilize Ndc80 levels through creating an alternative cellular state in meiotic prophase.

Next, we tested whether Aurora B/Ipl1 activity is disrupted in the *ama1∆* mutant, as Ipl1 downregulation would lead to Ndc80 stabilization. We found that the wild type and *ama1∆* cells had comparable levels of serine 10 phosphorylated histone H3 (Figure 6E), which is an established Aurora B/Ipl1 substrate (Hsu et al. 2000). Thus, Ipl1 activity is unaffected by Ama1 deletion. These observations led us to conclude that APC^Ama1^ acts downstream of Ipl1-dependent phosphorylation to mediate Ndc80 degradation.

### Defects in Ndc80 degradation predispose meiotic cells to chromosome segregation errors

How does Ndc80 degradation impact kinetochore activity and function? We used the *∆2-28* mutant to address this question. We first tested if this mutant had growth defects at higher temperature (37 °C) or on plates containing the microtubule depolymerizing drug benomyl. These two phenotypes are often exhibited by mutants with defects in kinetochore function (Spencer et al. 1990; Wigge et al. 1998; Hyland et al. 1999) and spindle assembly checkpoint (e.g. *mad2∆*) (Li and Murray 1991). We found the *∆2-28* mutant grew similarly as the wild type cells in both conditions (Figure 7A), demonstrating that these amino acids in Ndc80 are dispensable for normal kinetochore function in mitotic cells.

**Figure 7.**
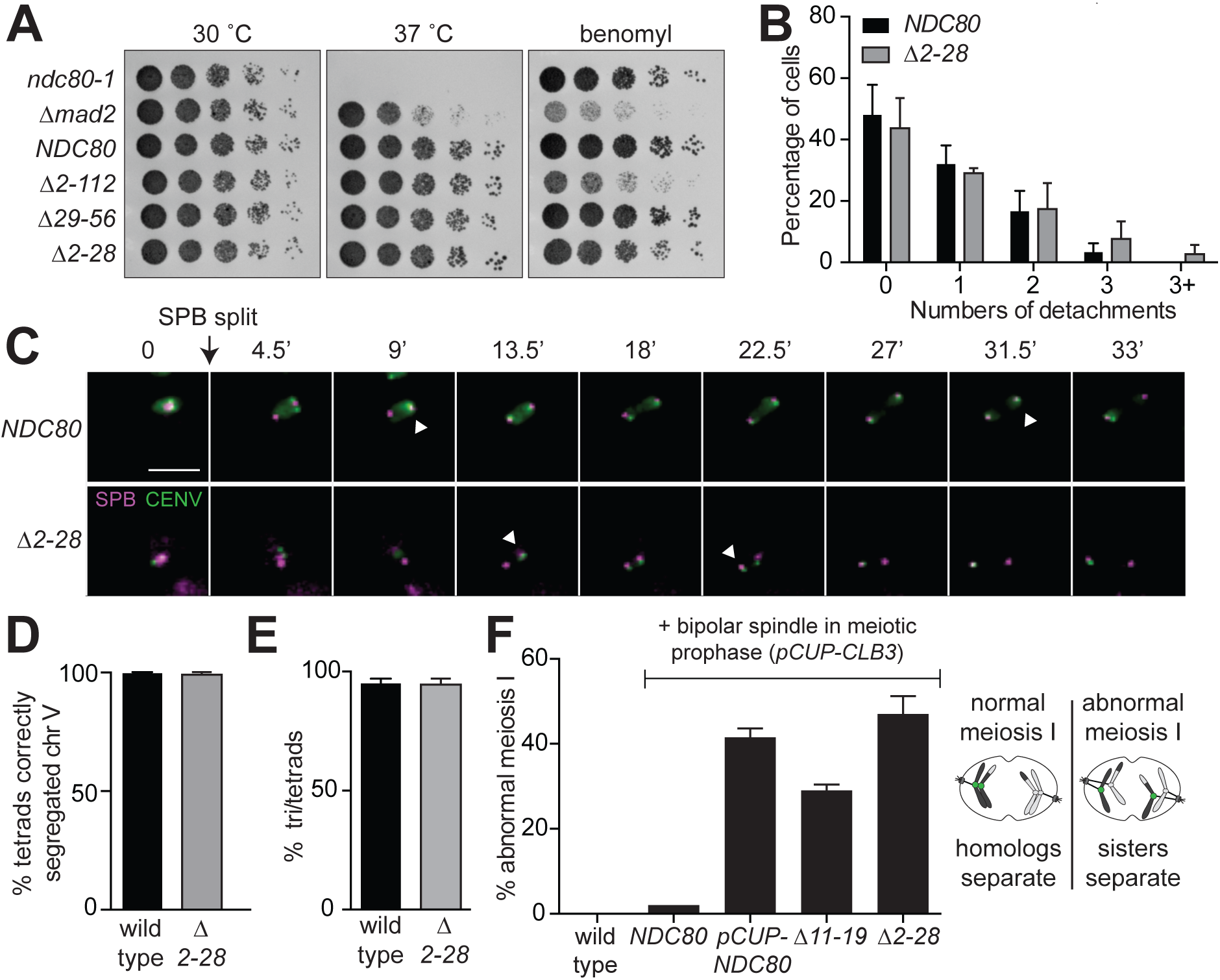
Ndc80 degradation regulates kinetochore composition in meiosis. (A) Growth phenotype associated with the various truncations of the Ndc80 N-terminal tail. Tested strains included the temperature-sensitive *NDC80* allele (*ndc80-1*, UB494), the spindle assembly checkpoint mutant *∆mad2* (UB700), wild type *NDC80* (UB3262), as well as the *NDC80* mutants that carry the following deletion: residues 2-112 (*∆2-112*, UB3275), residues 29-56 (*∆29-56)*, UB4695), and residues 2-28 (*∆2-28)*, UB5015). Cells were serially diluted and grown on plates containing nutrient rich medium (YPD) at 30 °C or 37 °C, as well as on a benomyl plate (15 µg/ml) at 23 °C. (b) Frequency distribution of the cells with a given number of kinetochore detachments and re-attachments for the experiment in (C). The mean percentage and the standard deviation for 3 biological replicates are displayed. (C) Kinetochore attachments and detachments for the *NDC80* (UB15905) and *∆2-28* (UB15873) strains in the *spo11∆* background. Each strain carries the TetR-GFP fusion protein, and both homologs of chromosome V have the centromeric *TetO* array (homozygous *CENV-GFP*, marked in green). Magenta: Spc42 fused with mCherry, which marks the spindle pole body (SPB). White arrow marks the kinetochore that would undergo detachment. The time when the SPBs were split is marked as 0. Scale bar: 5 µm. (D) The percentage of cells correctly segregated chromosome V for the wild type (UB21757) and *∆2-28* (UB21758) strains. These strains carry the TetR-GFP fusion proteins, and both homologs of chromosome V are marked by the centromeric *TetO* repeats (*CENV-GFP*). Strains were sporulated for 7 h before formaldehyde fixation. At least 90 cells were counted for each strain. For panel D and E, the mean and the range of 2 biological replicates are graphed. (E) Spore formation for the wild type (UB21756) and *∆2-28* (UB21758) strains. (F) Sister chromatid segregation in wild type (UB2942), *pCUP-CLB3 NDC80* (UB877), *pCUP-CLB3 pCUP-NDC80* (UB880), *pCUP-CLB3 NDC80(∆11-19)* (UB16561), and *pCUP-CLB3 NDC80(∆2-28)* (UB16565). All strains carry *pGAL-NDT80* and *GAL4.ER*, which allows reversible arrest in meiotic prophase. A pair of the chromosome V sister chromatids is marked by the centromeric *TetO* repeats. A schematic for normal and abnormal meiosis I is shown on the right. In an abnormal meiosis I, two separated GFP dots were observed in binucleates, one in each nucleus. Cells were sporulated for 5 h before addition of CuSO_4_ to induce cyclin Clb3 expression. Immediately after the induction, cells were released from the prophase arrest using ß-estradiol, which induces *NDT80*. Samples were taken 1 h 45 min after the release. The average fraction of binucleates that displayed sister segregation in meiosis I, as well as the range of 2 biological replicates, is graphed. 100 cells were counted per strain, per experiment.

To examine if Ndc80 degradation affects kinetochore function in meiosis, we monitored chromosome movements using the centromeric TetR/TetO GFP dot assay (Michaelis et al. 1997), where both homologs of chromosome V contained the TetO arrays that were integrated near the centromere and were bound by the TetR-GFP fusion protein (CENV-GFP). We performed our experiment in the *spo11∆* background because (1) these cells have long spindles that help resolve chromosome movements (Shonn et al. 2000) and (2) the homologous chromosomes fail to recombine (Klapholz et al. 1985), which renders the *spo11∆* cells unable to establish bi-orientation. As a result, multiple rounds of microtubule–kinetochore attachments and detachments occur for an extended period, as part of the error correction mechanism. Ipl1 inactivation is known to cause defects in error correction, which is manifested by a reduced frequency of kinetochore detachments in the *spo11∆* cells (Meyer et al. 2013). We found a similar distribution of kinetochore detachments between the wild type and *∆2-28* cells (Figure 7B-C), suggesting that the frequency of error correction, and by inference, the kinetochore function is unaltered upon Ndc80 stabilization. Consistently, over 95% of the wild type and the *∆2-28* tetrads correctly segregated their chromosomes, as shown by tracking the segregation of the homozygous CENV-GFP dots (Figure 7D and S7A). Both the wild type and *∆2-28* cells had >95% sporulation efficiency (Figure 7E) and >96% spore viability (Figure S7B-C), a metric that would be lowered if chromosomes mis-segregated. These results suggest Ndc80 degradation is dispensable for correcting erroneous microtubule-kinetochore interactions in meiosis.

Instead, we found that Ndc80 degradation modulated kinetochore activity by changing kinetochore composition. Remodeling kinetochore composition is crucial for proper meiosis I. In meiotic prophase, the outer kinetochore dissociates from the inner kinetochore to downregulate kinetochore activity, which prevents premature microtubule-kinetochore interactions and is crucial to establish a meiosis I-specific chromosome segregation pattern (Asakawa et al. 2005; Miller et al. 2012; Meyer et al. 2015; Chen et al. 2017). When the *∆2-28* or *∆11-19* mutant were combined with a mutant that prematurely forms spindle microtubules in meiotic prophase (*CUP-CLB3*), we observed that sister chromatids, instead of homologous chromosomes, segregated in meiosis I (Figure 7E). The same phenotype occurs when Ndc80 is overexpressed in meiotic prophase or when the LUTI-based repression of Ndc80 synthesis is disrupted (Miller et al. 2012; Chen et al. 2017). Therefore, the lack of Ndc80 turnover in meiotic prophase prematurely activates the kinetochore by providing a high abundance of Ndc80, which is the limiting subunit of the kinetochore activity in meiotic prophase, and predisposes cells to meiotic chromosome segregation errors.

## DISCUSSION

In this study, we have shown that Ndc80 degradation in meiosis is a temporally regulated process. The proteolysis of Ndc80 in meiotic prophase is independent of major chromosome remodeling events, but requires the ubiquitin ligase APC^Ama1^ and proteasome activity. Furthermore, Ndc80 degradation is coupled to Aurora B/Ipl1-mediated phosphorylation of Ndc80, a post-translational modification known for correcting erroneous microtubule–kinetochore attachments. The N-terminus of Ndc80, which includes a 27-residue sequence and the Aurora B/Ipl1 consensus sites, is both necessary and sufficient to drive the degradation of kinetochore proteins. The failure to degrade Ndc80 in meiotic prophase alters the kinetics of kinetochore dispersion, causes premature activation of kinetochores, and predisposes meiotic cells to chromosome segregation defects. All of these observations highlight the importance of regulating Ndc80 turnover in meiosis.

### Phosphorylation-mediated Ndc80 degradation in meiotic prophase

Multiple lines of evidence support a model in which Aurora B/Ipl1–dependent phosphorylation of Ndc80 triggers its degradation in meiotic prophase. First, Ipl1 is required for Ndc80 degradation. Second, phosphorylation of an Ipl1 consensus site on Ndc80 is detected by mass spectrometry in meiotic prophase, a stage when Ndc80 degradation occurs. Third, mutating the seven Ipl1 consensus sites leads to Ndc80 stabilization in meiotic prophase, suggesting that these sites are necessary for Ndc80 degradation. Altogether, these results are consistent with a phosphorylation-dependent degradation mechanism.

Given that Ndc80 phosphorylation is not disrupted when the 2-28 residues are deleted, we posit that this 2-28 segment may act in parallel or downstream of the Ipl1-dependent phosphorylation to promote Ndc80 degradation. These residues may serve as the binding site for protein factors that mediate Ndc80 proteolysis. It is possible that such factors are recruited by Ndc80 phosphorylation, or that phosphorylation alters the local conformation around the 2-28 residues to expose this segment for protein factor binding.

Our data are consistent with the idea that the ubiquitin ligase APC^Ama1^ and the proteasome act downstream of the Ipl1-dependent phosphorylation to mediate Ndc80 proteolysis. We excluded the possibility that the *ama1∆* mutation stabilizes Ndc80 by increasing cyclin/CDK activity or by downregulating Ipl1 activity (Figure 6C-E). It is currently unknown whether APC^Ama1^ directly ubiquitinate Ndc80 or regulate other player(s) in the degradation pathway to promote Ndc80 turnover. Interestingly, the APC/C degron repository predicts that Ndc80 has 4 D-box motifs and 3 KEN motifs, which are known recognition sites for the APC/C (Glotzer et al. 1991). However, these sites are located outside of the 112-residue N-terminal region, which is both necessary and sufficient for Ndc80 degradation in meiotic prophase. Furthermore, mutating the 4 D-boxes does not block Ndc80 degradation (Figure S7D). The D-box motif is also not required for the APC^Ama1^-dependent proteolysis of Ssp1 (Maier et al. 2007; Diamond et al. 2009). Future work to characterize the substrates of APC^Ama1^, as well as the mechanism of substrate recognition, will be important to fully grasp the meiotic regulation of Ndc80 turnover.

### Interplay between error correction and Ndc80 degradation

It is well established that Ndc80 phosphorylation by Aurora B/Ipl1 helps correct microtubule–kinetochore attachments that do not result in tension (Biggins 2013). Ndc80 phosphorylation is thought to weaken microtubule binding, resulting in microtubule detachment and thus providing the opportunity for the kinetochore to re-attach in the correct orientation. How is the error correction process interrelated with Ndc80 degradation given that both processes require Ndc80 phosphorylation by Aurora B? We found that disrupting Ndc80 degradation alone (*∆2-28*) does not appear to affect meiotic chromosome segregation (Figure 7B-E, S7A-C). This observation suggests that Ndc80 degradation is not required for error correction.

Instead, we propose that the completion of error correction may repress Ndc80 degradation in metaphase I, a stage when Ndc80 is stable (Figure 1E, S1B-C). Once the chromosomes attach to spindles properly, Ndc80 phosphorylation is removed by phosphatases to stabilize the microtubule attachments (Biggins 2013). Thus, it is possible that the Ndc80 levels become stable after the removal of Ndc80 phosphorylation in metaphase I. We note that it is unlikely that microtubule binding directly prevent Ndc80 degradation through steric effect because, at least in meiotic prophase, microtubule depolymerization cannot destabilize Ndc80 in *ipl1-mn* cells.

Our model presents a puzzle, however. As the major microtubule-binding site, Ndc80 is required to build new attachments during error correction. And yet, in our model, Ndc80 phosphorylation would cause its degradation at a time when the presence of Ndc80 is essential. This problem could be reconciled by the fact that Ndc80 protein is upregulated as cells exit from meiotic prophase (Chen et al. 2017), which may provide extra proteins to compensate for the loss due to degradation. In addition, the activity of the ubiquitin ligase APC^Ama1^ is repressed by Clb-CDK activity, which rises in prometaphase I (Oelschlaegel et al. 2005; Tsuchiya et al. 2011). Since APC^Ama1^ is required for Ndc80 degradation, the degradation mechanism of Ndc80 could be turned off after meiotic prophase, even before error correction is completed.

### Protein turnover as a mechanism to create meiosis-specific kinetochores

In vegetative growth, the yeast kinetochores transiently disassemble during DNA replication when the centromeric DNA replicates, but they remain bound to centromeres for the rest of the cell cycle. It has been shown that the subunit stoichiometry of the kinetochore is regulated during the cell cycle (Dhatchinamoorthy et al. 2017; Dhatchinamoorthy et al. 2019). However, it is unclear whether the protein levels of the kinetochore subunits are regulated in vegetative growth. Degradation of a few kinetochore proteins has been reported in yeast. These include Dsn1 (Akiyoshi et al. 2013), Cbf13 (Kaplan et al. 1997), and the centromeric histone Cse4 in budding yeast (Herrero and Thorpe 2016; Ohkuni et al. 2016; Cheng et al. 2017), as well as Spc7 in fission yeast (Kriegenburg et al. 2014). The degradation pathways of these subunits have been proposed to be quality controls that remove nonfunctional or excess proteins, rather than direct means to alter kinetochore function.

In contrast, regulating protein abundance is the key mechanism of controlling kinetochore activity in meiosis. The outer kinetochore disassembles in meiotic prophase and reassembles in prometaphase I. This disassembly is triggered by a reduction in Ndc80 levels, which occurs as a result of two separate mechanisms: LUTI-based repression of new Ndc80 synthesis and degradation of the existing Ndc80 pool. In meiosis, Ndc80 degradation is not a passive consequence of DNA replication but a targeted process mediated by Aurora B/Ipl1, which has also been implicated in preventing untimely spindle formation in meiotic prophase (Shirk et al. 2011; Kim et al. 2013). We propose that Ipl1 prevents premature microtubule–kinetochore interactions through two independent pathways: one by altering microtubule behavior, and the other by triggering Ndc80 proteolysis. This dual mechanism ensures that the kinetochore interacts with spindles only after meiotic prophase. This delay in microtubule– kinetochore interaction is required for proper meiotic chromosome segregation (Miller et al. 2012).

### Cells customize kinetochore activity by controlling kinetochore assembly or disassembly

Examples from various organisms indicate that regulating kinetochore assembly or disassembly is a common way to customize kinetochore activity based on the specific cell cycle stage or cell type. During mitotic exit in human cells, the outer kinetochore components, known as the KMN network, dissociate from the inner kinetochore in every cell cycle. A failure to do so causes chromosome mis-segregation in the next cell cycle (Gascoigne and Cheeseman 2013). In *C. elegans* oocytes, kinetochores facilitate homolog bi-orientation and error sensing in meiosis I, but they do not segregate the chromosomes. Instead, the ring-complex and microtubule motors, such as dynein, segregate the chromosomes, while the kinetochsore proteins disappear from the chromosomes in anaphase I (Dumont et al. 2010; Muscat et al. 2015; Davis-Roca et al. 2017; Laband et al. 2017). Lastly, in *C. elegans*, *Drosophila*, and iPS-derived human motor neurons, some or all of the core kinetochore proteins localize to the regions of synaptic neuropil and axons that are devoid of nuclei. There, they control axon outgrowth and dendritic extension in the developing sensory nervous system (Cheerambathur et al. 2019; Zhao et al. 2019). All these examples highlight that the timing and location of kinetochore assembly are tuned to the cell cycle and cell type.

Based on our finding that the Aurora B kinase regulates the turnover of the kinetochore subunit Ndc80, it will be interesting to test if other kinetochore restructuring events also depend on Aurora B. For example, the protein levels of Ndc80/Hec1 decline at the mitotic exit in human cells (Ferretti et al. 2010), but the mechanism is unknown. We posit that Aurora B and/or Aurora A may trigger Ndc80/Hec1 degradation at the mitotic exit in human cells, contributing to the KMN dissociation. In addition, the *C. elegans* Aurora B kinase, AIR-2, localizes to spindle microtubules during anaphase I (Schumacher et al. 1998; Speliotes et al. 2000; Romano et al. 2003). Thus, it is possible that AIR-2 phosphorylation causes the dissociation or degradation of one or more kinetochore subunits, leading to kinetochore disassembly at that time. Our discovery that Aurora B phosphorylation can cause the turnover of kinetochore proteins provides a new means by which Aurora B can influence kinetochore composition and activity: Aurora B not only regulates the construction but also the destruction of the kinetochore.

## MATERIALS and METHODS

### Yeast strains and plasmids

All strains used in this study were derivatives of SK1 unless specified. The strain genotypes are listed in Supplemental file 1. The centromeric TetR/TetO GFP dot assay was first described in Michaelis et al. 1997, the *ndc80-1* temperature-sensitive mutant in Wigge et al. 1998, *pCUP-NDC80* and *pCUP-CLB3* in Lee and Amon 2003; Miller et al. 2012, the meiotic-depletion alleles *pCLB2-3HA-CDC20* in Lee and Amon 2003, *pSCC1-3HA-CDC6* in Hochwagen et al. 2005, and *SPC24-6HIS-3FLAG* in Miller et al. 2016. The following alleles were generated at the endogenous gene loci using PCR-based methods (Longtine et al. 1998): *NDC80-3V5*, *pSCC1-3HA-IPL1*, *ndt80∆, spo11∆*, *pdr5∆*, *mad2∆*, *mad3∆*, *ama1∆*, *pCLB2-3HA-NDD1*, *clb4∆*, *NDC80-eGFP*, *MTW1-mCherry*, and *∆NDC80^LUTI^* (deletion of the 400 bp to 600 bp upstream of the *NDC80* coding region). The V5 tagging plasmid was kindly provided by Vincent Guacci (University of California, Berkeley).

Previously, an *NDC80* single integration plasmid was generated (Chen et al. 2017). This plasmid includes the SK1 genomic sequence spanning from 1000 bp upstream to 357 bp downstream of the *NDC80* coding region, as well as a C-terminal fusion of the 3V5 epitope to *NDC80* referred to as *NDC80-3V5* (Chen et al. 2017). All the single integration plasmids used in this study were derived from this *NDC80-3V5* parent vector.

We cloned the following mutant *NDC80* alleles by Gibson assembly (Gibson et al. 2009) using gBlocks gene fragments (IDT, Redwood City, CA)*: NDC80-7A-3V5*, *NDC80-14A-3V5*, *NDC80(4H to A)-3V5*, *NDC80(4H to L)-3V5, NDC80-11A-3V5*, and *NDC80-4D-box-3V5.* The *8lexO-LUTI-NDC80-3V5* construct, also cloned by Gibson assembly, has 8 *lex* operators (*lexO*) and the promoter from *CYC1* inserted at 536 bp upstream of the translation start site of *NDC80 ORF*. This insertion allows lexA-B112-ER to control the expression of *NDC80*^*LUTI*^ mRNA in a dose-dependent manner (Chia et al. 2017). The *8lexO-LUTI-mse-NDC80-3V5* construct has a mutation in the Ndt80 binding site (*mse*), which diminishes the Ndt80-dependent expression of *NDC80*^*ORF*^ mRNA (Chen et al. 2017). The *pNDC80-AME1-3V5* and *pNDC80-AME1-tail-3V5* (referred to as *AME1-tail* in the Results section) constructs are controlled by the *NDC80* promoter (1000 bp upstream of the *NDC80* coding region), which is sufficient to repress protein synthesis of the coding region in meiotic prophase (Chen et al. 2017). For the *pNDC80-AME1-tail-3V5* construct, the 2-112 residues of Ndc80 were fused to the C-terminus of Ame1.

In addition, systematic deletions of the N-terminal residues of Ndc80 (2-28, 29-56, 2-10, 11-19, 20-28, 57-112) were made on the *NDC80-3V5* parent vector using the Q5 Site-Directed Mutagenesis Kit (NEB, Ipswich, MA). All the single integration plasmids described above were digested with PmeI, transformed into yeast, and selected for integration at the *LEU2* locus.

To generate the *∆2-28*, *∆11-19*, and *NDC80-7A* alleles at the endogenous locus of *NDC80*, the CRISPR/Cas9 method was used (Anand et al. 2017). Oligonucleotides encoding the guide RNA (5’-TACATCACATGGACCCTCATCGG-3’) were cloned into a centromeric plasmid carrying a *URA3* maker and pPGK1-Cas9 (a gift from Gavin Schlissel, University of California, Berkeley, CA). This plasmid was co-transformed into yeast along with the repair templates amplified from the respective *LEU2* single integration plasmids by PCR (*∆2-28*, *∆11-19*, and *NDC80-7A* plasmids). For *NDC80-7A*, the following primers were used to amplify the repair template while introducing synonomous mutations to the PAM sequence to prevent re-editing:

(5’ACATGTGCTACATCACATGGACCCTCATCGGTTTGCTTCaCAAATACCAACTGCA ACATC-3’)

(5’CTCTTGAATAGCGCTTTGGAAGTTTTTGTCTCTTAGTGGtCTTGGATCTCTATTGC TCAG-3’)

The lowercase letters correspond to the synonymous mutations that abolish the PAM site: a C-to-A mutation at the 66^th^ nucleotide from the translation start site of Ndc80, as well as a G-to-A at the 351^st^ nucleotide.

### Yeast growth condition, synchronous sporulation, and media

To prepare for sporulation, diploid cells were grown in YPD (1% yeast extract, 2% peptone, 2% glucose, and supplemented with 22.4 mg/L uracil and 80 mg/L tryptophan) for 20-24 hours at room temperature or 30°C. For optimal aeration, the total volume of the flask exceeded the volume of the medium by 10-fold. Subsequently, cells were transferred to BYTA (1% yeast extract, 2% bacto tryptone, 1% potassium acetate, 50 mM potassium phthalate) at 0.25-0.3 OD_600_ cells/ml and grown for another 15-17 hours at 30°C. The cells were then pelleted, washed with sterile milliQ water, and resuspended at 1.85 OD_600_ cells/ml in sporulation (SPO) media (0.5% (w/v) potassium acetate [pH 7], 0.02% (w/v) raffinose) at 30°C unless specified.

Two methods were used to synchronize sporulation in this study. The *pCUP1-IME1 pCUP1-IME4* method was described in (Berchowitz et al. 2013), in which the endogenous promoters of *IME1* and *IME4* were replaced with the inducible *CUP1* promoter. To initiate synchronous sporulation, the expression of *IME1* and *IME4* was induced by adding 50 µM copper (II) sulphate 2h after cells were transferred to SPO. The *pGAL-NDT80 GAL4-ER* system was used to allow cells to undergo synchronized meiotic divisions (Carlile and Amon 2008). After incubating the sporulating diploids for 5-6 hours in SPO (see figure legend for the specific arrest duration for each experiment), cells were released from pachytene arrest by inducing *NDT80* expression with the addition of 1 µM β-estradiol.

### RNA extraction and RT-qPCR

A previously described RNA extraction protocol was modified as described below (Koster and Timmers 2015). RNA was extracted with acid phenol:chloroform:isoamyl alcohol (125:24:1; pH 4.7) and then isopropanol precipitated. Briefly, 1.85-3.7 OD_600_ of cells were harvested by centrifugation at 21000 rcf for 1 min, and frozen in liquid nitrogen. Pellets were thawed on ice, and vigorously shaken at 65°C in 400 µl TES buffer (10 mM Tris [pH 7.5], 10 mM EDTA, 0.5% SDS), 200 µl of acid-washed glass beads (Sigma), and 400 µl acid phenol for 30 min. The lysates were clarified by centrifugation at 21000 rcf at 4°C for 10 min. The aqueous phase was transferred to 300 µl chloroform, vortexed, and centrifuged at 21000 rcf at room temperature for 5 min. To precipitate the RNA, the aqueous phase was added to 400 µl isopropanol supplemented with 0.33 M NaOAc, incubated at −20°C for more than two hours, and centrifuged at 21000 rcf, 4°C, for 20 min. The precipitated RNA pellets were washed with 80% ethanol, air dried, and resuspended in nuclease-free water.

A total of 5 µg RNA was treated with DNase using the TURBO DNA-free kit (Thermo Fisher) and then reverse-transcribed into cDNA using Superscript II or II (Thermo Fisher) following the standard protocol. The single-stranded cDNA was quantified by 7500 FAST Real-Time PCR machine (Thermo Fisher) using SYBR green mix (Thermo Fisher). The primers used for the RT-qPCR reactions are: *NDC80*^*LUTI*^ (GGTTGAGAGCCCCGTTAAGT and TTGGCACTTTCAGTATGGGT); *PFY1* (ACGGTAGACATGATGCTGAGG and ACGGTTGGTGGATAATGAGC). The Ct value for each sample was determined by taking the average Ct of triplicate reactions, and the levels of the *NDC80*^*LUTI*^ transcript were normalized to those of the *PFY1* transcript.

### Single molecule RNA fluorescence in situ hybridization (smFISH)

Single-molecule RNA FISH was performed exactly as described previously (Chen et al. 2017). To quantify smFISH spots, the maximum-intensity projection of z-stacks was generated in FIJI (Schindelin et al. 2012), different channels were split, and the images were analyzed by the StarSearch Program (Raj et al. 2008). Cell boundaries were hand-drawn. The thresholds were adjusted such that they fell within “plateaus,” as previously described (Raj et al. 2008). The same threshold was used for a given probe set in each experimental replicate. An *NDC80*^*LUTI*^ transcript was identified as a colocalized spot where the two spots found in either channel have pixels overlapped for over 50%. The number of *NDC80*^*ORF*^ transcripts in each cell was calculated by subtracting the number of *NDC80*^*LUTI*^ spots from the total number of spots with *NDC80*^*ORF*^ signal. To compare between different strains, data were pooled from two biological replicates, and the non-parametric Wilcoxon Rank Sum test was performed with Prism 6 (GraphPad). The relative frequency histograms were normalized so that the maximum bin height is the same across all the histograms in this study.

### Fluorescence microscopy

To fix cells, formaldehyde was added to ~1 OD_600_ cells to a final concentration of 3.7%, incubated at room temperature for 15 min, washed once with KPi/sorbitol buffer (100 mM potassium phosphate [pH 7.5], 1.2 M sorbitol), and resuspended in the VectaShield Antifade Mounting Medium with DAPI overnight before imaging (Vector Labs, Burlingame, CA). The antifade medium preserved the samples from photobleaching during imaging. All the images were acquired as z-stacks of 20 slices (0.3 µm step size) using a DeltaVision Elite wide-field fluorescence microscope, a PCO Edge sCMOS camera (Delta Vision, GE Healthcare, Sunnyvale, CA), a 100x/1.40 oil-immersion Plan Apochromat objective, and the following filters: DAPI (EX390/18, EM 435/48), GFP/FITC (EX475/28, EM525/48), and mCherry (EX575/25, EM625/45). Images were deconvolved in softWoRx software (softWoRx, *GE Healthcare*, *USA*) with a 3D iterative constrained deconvolution algorithm (enhanced ratio) with 15 iterations.

### Kinetochore classification and quantification

The three classes of kinetochores were classified as follows: The “clustered” class was defined by the presence of a single Mtw1-mCherry focus in a cell, or the presence of less than 3 distinct Mtw1-mCherry foci. For this class, Ndc80 signal was always detected and co-localized with the Mtw1-mCherry signal in the cells, and the DAPI signal was often compacted and round, without thread-like structures. The “dispersed” class was defined by more than 6 distinct Mtw1-mCherry foci located throughout the projected DAPI area of the cell. For this class, the DAPI signal was almost always thread-like, suggesting a pachytene state. The “partially clustered” class was defined by the presence of at least 3 distinct Mtw1-mCherry foci and all foci were located within the same half of the projected DAPI boundary of the cell.

### Live-cell imaging

Time-lapse microscopy was performed in an environmental chamber heated to 30°C. Strains were induced to sporulate by transferring to SPO and halted in meiotic prophase using the *pGAL-NDT80 GAL4-ER* block until 5h in SPO. Cells were then released with 1 µM β-estradiol, and 30 min later, immobilized on concanavalin A-coated, glass-bottom 96-well plates (Corning) and cultured in 100 µl SPO (with 1 µM β-estradiol) medium in the pre-warmed environmental chamber. Z-stacks of 5 slices (1-µm step size) were acquired using a 60×/NA1.42 oil-immersion Plan Apochromat objective, mCherry filter (32% T, 25-ms exposure) and FITC filter (10% T, 25-ms exposure), at a frequency of 45 sec for 90 min.

The following pipeline was used to quantify kinetochore detachment events. First, the movie was maximum-projected using FIJI (Schindelin et al. 2012). Next, the time of spindle pole body (SPB) separation and each kinetochore detachment event were tracked. The time of SPB separation is defined as the first frame when two distinct mCherry foci were detected. Kinetochore attachment is defined as a CENV-GFP focus staying in the proximity of one SPB (less than a third of the pole-to-pole distance) for at least 3 consecutive time frames. A detachment event occurred when a CENV-GFP focus switched SPB or crossed the midline between the two poles (the CENV-GFP focus often stayed in the midzone for more than 3 consecutive frames before being recaptured by the original SPB). The total number of detachment events occurred after the splitting of the SPBs was quantified for each cell that maintained in the focal plane for the entire movie. The percentage of cells that underwent a given number of detachment events was calculated and graphed.

### Denaturing immunoprecipitation (IP) and mass spectrometry (MS)

Denatured protein extracts were prepared as described in Sawyer et al. 2019, with the following modifications. After cells were harvested and dried completely, 150 µl of zirconia beads and 150 µl of lysis buffer (50 mM Tris [pH 7.5], 1 mM EDTA, 2.75 mM DTT, 1x PhosSTOP (Roche), 0.48 µg/ml Pefabloc SC (Sigma), 2 mM pepstatin A, and 2x cOmplete EDTA Free (Roche)) were added to each tube. Cells were lysed mechanically on a Mini-Beadbeater-96 (BioSpec Products) at room temperature for 5 min. SDS was added to a final concentration of 1%, the extracts were boiled at 95°C for 5 min, and NP-40 lysis buffer (50 mM Tris [pH 7.5], 150 mM NaCl, 1% NP-40, and 5% glycerol, 1x PhosSTOP, 0.64 µg/ml Pefabloc SC (Sigma), 2.67 mM pepstatin A and 3x cOmplete EDTA Free) was added to a final volume of 1.5 ml (i.e., diluting SDS to 0.1%). Lysates were clarified by centrifuging at 15000 rcf, 4°C, for 15 min. The total cleared lysate for each sample was pooled from 3 tubes (3,825 µl total) and added to 80 µl pre-equilibrated anti-V5 agarose beads (Sigma), and incubated in loBind tubes (Eppendorf, Hamburg, Germany) at 4°C for 2h with rotation. Using loBind tubes and low-adhesion tubes (USA Scientific, Orlando, Florida) improved the amount of protein recovered from the IP. In each MS experiment, the cleared lysates were pooled from 6 tubes for each sample, therefore two IP reactions were set up in parallel. The beads of the parallel IP reactions were washed once with the 3 ml High Salt Wash buffer (50 mM Tris, pH 7.5, 0.5 M NaCl, 1 mM EDTA, and 1% NP-40), combined into one low-adhesion tube (i.e., 160 µl beads final), and then washed once with 950 µl of High Salt Wash buffer. Next, the beads were washed twice with each of the following: (1) 950 µl Buffer 2 (50 mM Tris [pH 7.5], 150 mM NaCl, 10 mM MgCl_2_, 0.05% NP-40, and 5% glycerol); and (2) 950 µl Buffer 3 (50 mM Tris [pH 7.5], 150 mM NaCl, 10 mM MgCl_2_, and 5% glycerol). After the last wash, the wash buffer was aspirated completely, and the beads were resuspended in 80 µl trypsin buffer (2 M Urea, 50 mM Tris [pH 7.5], 5 µg/ml trypsin) to digest the bound proteins at 37°C for 1h with agitation. The beads were centrifuged at 100 rcf for 30 sec, and the digested peptides (the supernatant) were collected. The beads were then washed twice with 60 µl Urea buffer (2 M Urea, 50 mM Tris [pH 7.5]). The supernatant of both washes was collected and combined with the digested peptides (i.e. the final volume is 200 µl). After brief centrifugation, the combined digested peptides were cleared from residual beads and frozen in liquid nitrogen.

MS was performed by the Vincent J. Coates Proteomics/Mass Spectrometry Laboratory at the University of California, Berkeley, as described in (Sawyer et al. 2019). The mudPIT method was used. A nano LC column (packed in a 100-µm inner-diameter glass capillary with an emitter tip) is composed of 10 cm of Polaris c185 µm packing material (Varian), followed by 4 cm of Partisphere 5 SCX (Whatman). The column was loaded with a pressure bomb, washed extensively with buffer A, and then directly coupled to an electrospray ionization source mounted on a LTQ XL linear ion trap mass spectrometer (Thermo Fisher). Chromatography was performed on an Agilent 1200 HPLC equipped with a split line to deliver a flow rate of 300 nl/min. Peptides were eluted using either a four-step or an eight-step mudPIT procedure (Washburn et al. 2001). Buffer A: 5% acetonitrile and 0.02% heptafluorobutyric acid (HFBA). Buffer B: 80% acetonitrile and 0.02% HFBA. Buffer C: 250 mM ammonium acetate, 5% acetonitrile, and 0.02% HFBA. Buffer D: the same as buffer C, but with 500 mM ammonium acetate. Proteins were identified with the Integrated Proteomics Pipeline software (IP2; Integrated Proteomics Applications) using the parameters and cutoffs exactly as described by (Sawyer et al. 2019).

### Native IP, Co-IP, TMT quantitative MS

For the TMT quantitative MS and co-IP experiments, cells were sporulated in 2% SPO (2% (w/v) potassium acetate [pH 7], 0.02% (w/v) raffinose, supplemented with 40 mg/l uracil, 20 mg/l histidine, 20 mg/l leucine, 20 mg/l tryptophan) at 30°C. Increasing the concentration of acetate and supplementing amino acids improved sporulation efficiency. Four hours after the cells were transferred to 2% SPO, 200 mM PMSF (dissolved in ethanol) was added to cultures to a final concentration of 2 mM. For the MS and the co-IP experiments, 462 or 92 OD_600_ of cells, respectively, were harvested by centrifuging at 1700 rcf, 4°C, for 2 min. The supernatant was discarded, and the cell pellets were transferred to 2-ml tubes, centrifuged at 21000 rcf, 4°C, for 1 min. The supernatant was aspirated completely, and the pellets were snap frozen in liquid nitrogen.

To extract proteins, pellets were thawed on ice, and 500 µl of NP-40 Native IP Buffer was added (50 mM Tris [pH 7.5], 150 mM NaCl, 2 mM MgCl_2_, 1% NP-40, supplemented with PhosSTOP tablet (1 tablet used in 10 ml buffer, Sigma), cOmplete ULTRA protease inhibitor cocktail (1 tablet used in 20 ml buffer, Sigma), 0.64 µg/ml Pefabloc SC (Sigma), and 2.7 µM Pepstatin A). Cells were lysed with 500 µl of zirconia beads (BioSpec) in a Fast-Prep-24 5G (MP Biomedicals LLC, Irvine, CA) using the "*S. cerevisiae*" setting at room temperature for 40 s. Next, the cell lysates were chilled in an ice water bath for 2 min. The tubes were punctured with a G20.5 needle, and the lysates were collected in 15-ml tubes (precooled to −20°C) by centrifuging at 3000 rpm, 20-40 sec. Lysates were clarified by centrifugation at 21000 rcf, 4°C, for 20 min and snap frozen in liquid nitrogen. The protein concentration of each lysate was determined by Bradford Assay (Bio-rad, Hercules, CA).

Ndc80-3V5 IP was performed at a protein concentration of 10 mg/ml in NP-40 Native IP Buffer. For the TMT quantitative MS experiment, a total of 38-40 mg protein, pooled from 10 tubes of lysates, was added to 167 µl pre-equilibrated anti-V5 agarose beads (Sigma) in a total volume of 4 ml (i.e., beads constituted 1/24 of the total IP volume), incubated at 4°C for 2.5h with rotation, and then washed with each of the following: (1) once with 3.5 ml NP-40 Native IP Buffer, (2) once with 950 µl NP-40 Native IP Buffer, (3) twice with 950 µl Buffer 2, and (4) twice with 950 µl Buffer 3. After the last wash, the proteins on the beads were digested in trypsin buffer, washed with Urea buffer, and frozen in liquid nitrogen as described in the Denaturing IP section. The TMT quantitative MS was performed as described in (Cheng et al. 2018).

The co-IP experiments were performed similarly, except for the following changes: a total of 4 mg protein was incubated with 27 µl of anti-V5 agarose beads in a total volume of 400 µl, and washed twice with 950 µl NP-40 Native IP Buffer before washing with Buffer 2 and Buffer 3. After the last wash, the wash buffer was aspirated until 100 µl was left, and 50 µl 3x SDS sample buffer (187.5 mM Tris [pH 6.8], 6% mercaptoethanol, 30% glycerol, 9% SDS, 0.05% bromophenol blue) was added to the beads before boiling for 5 min. After brief centrifugation, the eluted proteins were separated by SDS-PAGE as described in the immunoblotting section.

### Purification of the Ndc80 complex

Native Ndc80 complex was purified from asynchronously growing *S. cerevisiae* cells (UB16284 and UB19957) through anti-Flag immunoprecipitation of Spc24-6His-3Flag as described in (Miller et al. 2016). Cells were grown in YPD. Protein lysates were prepared by lysing cells in a Freezer/Mill (SPEX SamplePrep) submerged in liquid nitrogen (Sarangapani et al. 2014). Lysed cells were resuspended in buffer H (BH) (25 mM HEPES [pH 8.0], 2 mM MgCl2, 0.1 mM EDTA, 0.5 mM EGTA, 0.1% NP-40, 15% glycerol, and 750 mM KCl), supplemented with the protease inhibitors (20 µg/ml leupeptin, 20 µg/ml pepstatin A, 20 µg/ml chymostatin, 200 µM phenylmethylsulfonyl fluoride) and the phosphatase inhibitors (0.1 mM Na-orthovanadate, 0.2 µM microcystin, 2 mM ß-glycerophosphate, 1 mM Na pyrophosphate, and 5 mM NaF). The resuspended cell lysates were ultracentrifuged at 98,500 g at 4°C for 90 min. Dynabeads conjugated with anti-Flag antibody (Sigma) were incubated with extract at 4°C for 3h with constant rotation, followed by three washes with BH containing the protease inhibitors, phosphatase inhibitors, 2 mM dithiothreitol (DTT), and 1 M KCl. The beads were further washed twice with BH containing 150 mM KCl and the protease inhibitors. Associated proteins were eluted from the beads by gently agitating the beads in elution buffer (0.5 mg/ml 3Flag peptide in BH with 150 mM KCl and the protease inhibitors) at room temperature for 30 min.

### *In Vitro* Kinase assay

The kinase assay was performed using a recombinant variant of Ipl1 (AurB*) that was created by fusing the C-terminal activation box of Sli15 to Ipl1 (a kind gift from S. Biggins’s lab, the Fred Hutchinson Cancer Research Center). The kinase-dead (KD) mutant of AurB* contains the K133R mutation. These two constructs were first reported in de Regt et al. 2018. To begin the reaction, in a volume of 6 µl, 6.63 ng of the purified WT-Ndc80 or Ndc80(∆2-28) was incubated with 1 µM AurB* or KD in BH0.15 (same ingredients as BH in the previous section except with 150 mM KCl) for 4 min at room temperature. Next, 6 µl of hot ATP mix (1.1 µl of 100 mM ATP, 3 µl of 6,000 Ci/mmol 10 mCi/ml g-32P ATP, and 275 µl of BH0.15) was added to the kinase/substrate mix. After 1 min or 5 min incubation at 25°C (see Figure Legend for details), reactions were stopped by adding 6 µl of 3x SDS sample buffer, and boiling at 95°C for 5-10 min. Proteins were separated by SDS-PAGE as described in the immunoblot section. The gel was fixed and silver stained based on the manufacturer’s instructions (Thermo Scientific). After the staining was stopped by incubating the gel in 5% acetic acid for 10 min, the gel was dried by vacuum at 80°C for 1h. The dried gel was exposed to a phosphor storage screen for over 24 hours. Screens were imaged with a Typhoon Scanner (GE Healthcare).

### Immunoblotting

To extract proteins from cells, ∼4 OD_600_ units of cells were treated with 5% trichloroacetic acid for at least 15 min at 4°C. After an acetone wash, the cell pellet was air dried, lysed with acid-washed glass beads (Sigma) in 100 µl of lysis buffer (50 mM Tris [pH 7.5], 1 mM EDTA, 2.75 mM DTT, cOmplete EDTA Free protease inhibitor cocktail (Roche, Basel, Switzerland)) using a Mini-Beadbeater-96 (Biospec Products). Next, 3x SDS sample buffer (187.5 mM Tris, pH 6.8, 6% 2-mercaptoethanol, 30% glycerol, 9% SDS, and 0.05% bromophenol blue) was added and the lysate was boiled for 5 min. Proteins were separated by SDS-PAGE using 4%-12% Bis-Tris Bolt gels (Thermo Fisher) and transferred onto nitrocellulose membranes (0.45 μm, Bio-rad, Hercules, CA) using a semi-dry transfer apparatus (Trans-Blot® Turbo^TM^ Transfer System, Bio-rad). The membranes were blocked for > 30 min with Odyssey® Blocking Buffer (PBS) (LI-COR Biosciences, Lincoln, NE), and incubated in primary antibodies overnight at 4°C. Membranes were washed with PBST (phosphate buffered saline with 0.01 % tween-20) and incubated with an anti-mouse secondary antibody conjugated to IRDye® 800CW at a 1:15,000 dilution (926-32212, LI-COR Biosciences) and/or an anti-rabbit antibody conjugated to IRDye® 680RD at a 1:15,000 dilution (926-68071, *LI-COR* Biosciences) for greater than 1h at room temperature. Immunoblots were imaged and quantified using the Odyssey® system (LI-COR Biosciences).

Antibodies used in this study: mouse anti-V5 antibody (R960-25, Thermo Fisher) used at a 1:2,000 dilution; rabbit anti-hexokinase antibody (H2035, US Biological, Salem, MA) used at a 1:20,000 dilution; rabbit anti-Sis1 antibody (kind gift from E. Craig) used at a 1:5000 dilution, and a rabbit anti-H3 serine 10 phosphorylation antibody (06-570, Sigma) used at 1:1000 dilution.

The levels of each protein of interest were typically normalized to Hxk2, except for in Figure 6F. For the quantification in Figure 6F, Ndc80 levels were not normalized to that of Hxk2 because Hxk2 levels declined after cycloheximide treatment for the *ama1∆* strain. To reflect the experimental variations, we performed 5 independent biological replicates and graphed the mean and the standard errors of the mean for all 5 replicates.

### Spot growth assay

Cells were grown on YPG (2% glycerol + YEP) plates overnight, resuspended in milliQ H_2_O, and then diluted to an OD_600_ of 0.1 or 0.2. 5-fold serial dilutions were performed. Cells were spotted onto YPD plates or the YPD plates supplemented with 15 µg/ml benomyl (benomyl plates). The cells on the YPD plates were incubated at 30°C for 1-2 days, and those on benomyl plates were incubated at 23°C for 2 days.

### Reproducibility

All the experiments in this study were performed at least two times independently, and one representative result was shown for all western blots. All the RT-qPCR experiments were performed three time independently, with the mean and standard errors of the mean graphed.

## ACKNOWLEDGEMENTS

We thank Michael Rape, David Drubin, Rebecca Heald, and all members of the Ünal and Brar labs for experimental suggestions and critiques of this manuscript. This work was supported by funds from the Pew Charitable Trusts (00027344), Damon Runyon Cancer Research Foundation (35-15) and National Institutes of Health (DP2 AG055946-01) to EÜ, and an NSF Graduate Research Fellowship Grant No. DGE-1106400 to JC.

## AUTHOR CONTRIBUTIONS

JC: conceptualization, data curation, formal analysis, investigation, methodology, validation, visualization, and drafted the manuscript. AL, ENP, and HL: investigation, formal analysis, validation, visualization, and manuscript editing. LAK: investigation, data curation, methodology, and formal analysis for the denaturing IP, as well as manuscript editing. RE: provided key resources (purifying Ndc80 complex) and manuscript editing. RMH: investigation for the truncation analysis of Ndc80 and manuscript editing. JKK: investigation and formal analysis for quantitative mass spectrometry. MJ: supervision and project administration. EÜ: conceptualization, formal analysis, methodology, visualization, supervision, project administration, funding acquisition, and drafted the manuscript.

